# The influence of the pre-membrane and envelope proteins on structure, pathogenicity and tropism of tick-borne encephalitis virus

**DOI:** 10.1101/2024.10.16.618642

**Authors:** Ebba Rosendal, Kyrylo Bisikalo, Stefanie M.A. Willekens, Marie Lindgren, Jiří Holoubek, Pavel Svoboda, Amanda Lappalainen, Ebba Könighofer, Ekaterina Mirgorodskaya, Rickard Nordén, Federico Morini, William Rosenbaum, Daniel Růžek, Ulf Ahlgren, Maria Anastasina, Andres Merits, Sarah J Butcher, Emma Nilsson, Anna K Överby

**Author notes:** Institute of Organic Chemistry and Biochemistry, Czech Academy of Sciences, Prague, Czech Republic (Pavel Svoboda).

## Abstract

Tick-borne encephalitis virus (TBEV) is a neurotropic flavivirus that causes thousands of human infections annually. Viral tropism in the brain is determined by the presence of necessary receptors, entry factors and the ability of the virus to overcome host defenses. The viral structural proteins, pre-membrane (prM) and envelope (E), play an important role in receptor binding, membrane fusion, particle maturation, and antibody neutralization. To understand how these proteins influence virus distribution and tropism in the brain, we generated a chimeric virus harboring the prM and ectodomain of E from TBEV in the background of the low pathogenic Langat virus (LGTV). We solved the atomic structures of both the chimeric virus and LGTV to compare them to the known TBEV structure. We show that this chimeric virus remains low-pathogenic, while being structurally and antigenically similar to TBEV. Using 3D optical whole brain imaging combined with immunohistochemistry, we found that both LGTV and the chimeric virus primarily infect cerebral cortex, with no significant differences in their localization or tropism. In contrast, TBEV shows high infection of the cerebellum and strong preference towards Purkinje cells, indicating that the non-structural proteins are important for determining TBEV tropism in the brain. Together, this provides new insights into the roles of the structural and non-structural proteins of tick-borne flaviviruses.

## Introduction

Tick-borne encephalitis virus (TBEV) is considered one of the most medically important arthropod-borne viruses in Eurasia and is estimated to result in 10,000-12,000 hospitalizations every year [1]. Symptoms following TBEV infection may range from mild flu-like symptoms to severe meningoencephalitis or encephalomyelitis with varying degrees of cognitive dysfunction [2]. TBEV is a member of the genus *Orthoflavivirus,* family *Flaviviridae*. These are enveloped viruses with a diameter of 50 nm and a ∼11 kb positive-sense RNA genome that contains a single open reading frame. This encodes a single polyprotein that is cleaved into 10 viral proteins, three of which are structural proteins that form the virion: the capsid (C), pre-membrane (prM), and envelope (E) proteins. The C protein binds to viral RNA to form the nucleocapsid (NC), and to lipids for NC budding [3]. The E protein is a class II fusion protein, and the largest of the structural proteins [4]. It is the main protein responsible for receptor binding and fusion, as well as the major target for neutralizing antibodies [5–9]. The prM protein is a precursor which is cleaved by the host cell protease furin into mature M protein and a pr peptide. This cleavage occurs in the trans-Golgi network (TGN) and is required for virus particle maturation [10].

Neurons are considered the main target cells for TBEV in the central nervous system (CNS) and primary neurons are highly susceptible to infection *in vitro* [11–13]. Understanding viral tropism in the brain is important as it is an underlying mechanism of pathology. It is determined by multiple factors, such as: the distribution of viral receptors, expression of antiviral host factors, the ability of the virus to prevent type I interferon (IFN) response, and the interplay between the adaptive immune system and brain resident cells [12, 14, 15]. Langat virus (LGTV) is a tick-isolate with naturally low pathogenicity. It has approximately 85% sequence similarity with TBEV on the amino acid level and is often used in research as a biosafety level 2 (BSL-2) model virus of more pathogenic tick-borne flaviviruses [14, 16, 17]. Recently, we used Optical Projection Tomography (OPT) in combination with higher resolution imaging techniques to visualize LGTV infection in whole mouse brain in 3D and showed that LGTV primarily infected neurons in the olfactory and somatosensory system in wild-type (WT) mice [18]. Meanwhile, a defective type I IFN response alters the global and cellular tropism of LGTV, and increases the susceptibility of choroid plexus, meninges and microglia in the cortex and the olfactory bulb [18]. However, it is not clear how well the tropism of LGTV corresponds to that of TBEV and which the viral factors are that influence tropism.

It has previously been established that exchanging the structural proteins between different flaviviruses is well tolerated, and chimeric TBEVs have been generated by inserting the prM and E proteins into various genetic backgrounds, including LGTV [19–21]. The resulting LGTV/TBEV chimeras were low-pathogenic but remained neuroinvasive and neurovirulent in animal models [20, 22]. Here, we investigated the influence of the structural proteins prM and E on virus structure, antigenicity, pathogenicity, virus distribution and tropism in the brain. We thoroughly characterized a chimeric virus containing the prM and ectodomain of E (ecto-E) proteins of TBEV in a LGTV genetic background. We show that it displays the pathogenicity profile of LGTV, is structurally coherent with both TBEV and LGTV and that its tropism in the brain overlaps significantly with LGTV.

## Results

### Chimeric LGTV^T:prME^ shows similar kinetics but increased viral spread *in vitro* compared to parental LGTV

We designed a chimeric virus based on the infectious clone of LGTV, replacing the prM protein and the ecto-E protein with those of TBEV strain 93/783 [23] (Figure 1A). Initial characterization of this chimeric virus (LGTV^T:prME^) *in vitro* showed that the kinetics of foci formation was similar to that of the recombinant parental LGTV, but slower than recombinant TBEV. However, the morphology of the foci was more similar to TBEV (Figure 1B). Viral replication and production of progeny virus following infection in A549 cells were also similar to LGTV, but lower than TBEV (Figure 1C-D). Next, we investigated the ability of chimeric virus to spread in cell culture, by infecting A549 cells at a low MOI (0.01) and quantifying the percentage of infected cells by immunostaining. We found that, at 48 hours post-infection (h.p.i.), TBEV showed the highest percentage of infected cells (27%), indicating faster kinetics than LGTV^T:prME^ (4%) and LGTV (1%) (Figure 1E). At 72 h.p.i., LGTV^T:prME^ showed the highest percentage of infected cells (72%), while for LGTV it remained low (11%), suggesting that the chimeric virus holds an advantage in cell-to-cell spread over the parental LGTV. The percentage of infected cells for TBEV did not increase between 48 and 72 h.p.i, most likely because of the high cytopathic effect of the virus observed at later time points (Figure 1F). Together, this indicates that exchanging prM and ecto-E of LGTV to those of TBEV does not influence virus replication efficiency nor the production of infectious particles, but slightly increases the virus’ ability to spread in cell culture.

**Figure 1.**
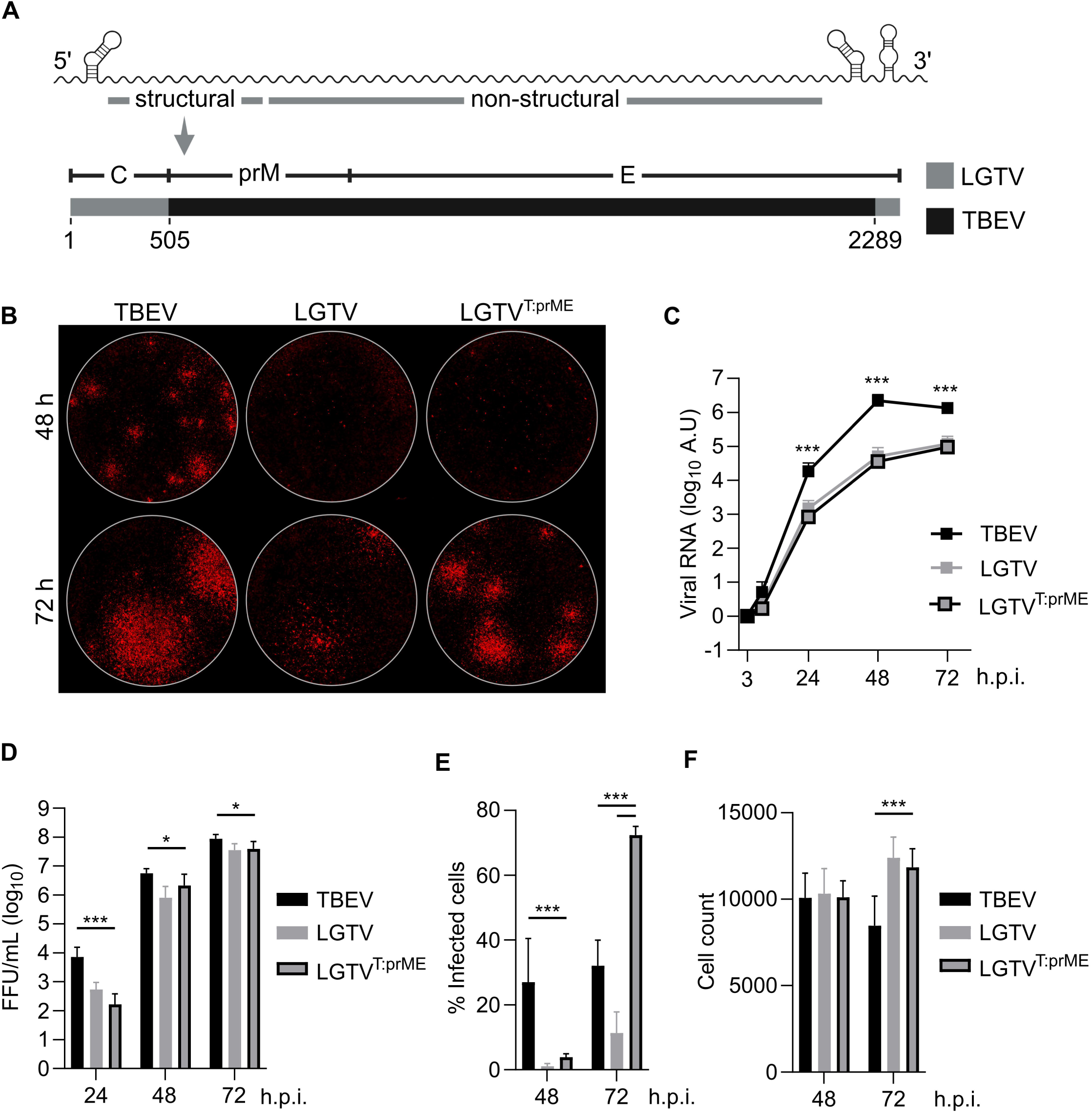
*In vitro* characterization of chimeric LGTV with structural proteins prM and E of TBEV (LGTV^T:prME^). **A)** Design of chimeric LGTV (LGTV^T:prME^). prM and ecto-E from TBEV strain 93/783 were used to replace their counterparts in LGTV strain TP21. Numbering represents positions of nucleotides in the nucleotide sequence of LGTV. **B)** Viral foci size and morphology of recombinant TBEV, LGTV and LGTV^T:prME^ on A549 *MAVS^-/-^* cells, 48 h or 72 h post-infection. Viral replication **C)** and virus release **D)** in A549 cells infected at MOI 0.1. Viral RNA levels were quantified using RT-qPCR and normalized using the ΔΔCt method to mRNA of actin at the 3 h time point (input control). Infectious virus in supernatant quantified as FFU/ml. Data from three independent experiments is shown as mean and SD; statistical significance calculated using Mann-Whitney test, * p < 0.05, ***p < 0.001. **E)** Viral spread in A549 cells infected at MOI 0.01. The percentage of infected cells was calculated by immunolabeling viral E and NS5 proteins and comparing the number of positive cells to the total cell count visualised using DAPI staining. **F)** Total cell count (DAPI) following infection. Data from three independent experiments performed in triplicates is shown as mean and SD; statistical significance calculated by parametric Student’s t test, *** p <0.001.

### Chimeric LGTV^T:prME^ is genetically stable in cell culture

Combining heterologous genes in chimeric viruses may cause mismatches in protein interactions resulting in accumulation of compensatory mutations over time. To assess this, we performed a 10x serial passaging of the chimeric virus and LGTV in cell culture followed by sequencing. We found that the chimeric virus showed a slightly lower number of total mutations (defined as >10% of sequencing reads) compared to LGTV, 15-20 and 32-33, respectively (Supplementary Table 1). Importantly, no missense mutation in the chimeric virus was observed in majority (>50%) of sequencing reads across all three replicates (Supplementary Table 2), suggesting that no consistent compensatory mutations arose during multiple passaging in cell culture.

### Chimeric LGTV^T:prME^ is structurally and antigenically similar to TBEV

We determined the structures of LGTV and chimeric LGTV^T:prME^ using cryogenic electron microscopy (cryo-EM) and three-dimensional image reconstruction (Supplementary Table 3). We established a rapid purification protocol using Capto™ Core 700 to remove contaminants from the VP-SFM culture medium, followed by concentration by ultrafiltration and vitrification. The electron micrographs of virus preparations contained mainly intact spherical virions with a size of ∼50 nm in diameter, along with some broken particles (Supplementary Figure 1A). We generated a 3.22 Å resolution map of LGTV^T:prME^ and a 3.82 Å resolution map of LGTV (Figure 2A-C, and Supplementary Figure 1A-D, Supplementary Table 3). We compared both atomic models to the previously published model of TBEV isolate Kuutsalo-14 [24]. The new models included two extra residues in the C-terminus of the E protein (residues 495 and 496) and the LGTV^T:prME^ model has one more residue in the C-terminus of the M protein (residue 75). Both structures are consistent with those of TBEV [8, 24], with the root mean square deviation (RMSD) of Cα atoms across protein chains below 0.7 Å for the E proteins and below 1.5 Å for the M proteins (Figure 2D, Supplementary Table 1). The ordered lipid head groups are also present in similar positions [24], hence, there are no major structural rearrangements in chimeric virus compared to the TBEV Kuutsalo-14 isolate. There are three occurrences of M in the asymmetric unit. In one of these three, we modelled two alternative conformations for residues 25-74 of LGTV^T:prME^. One conformation is identical to LGTV and TBEV, maintaining the pi-stacking interactions between M residues Trp37 at the raft interfaces, as well as the salt bridges between Lys40 and Glu33 of the respective chains, whereas the alternative conformation disrupts these tight interactions. (Figure 2E, Supplementary Figure 1E).

**Figure 2.**
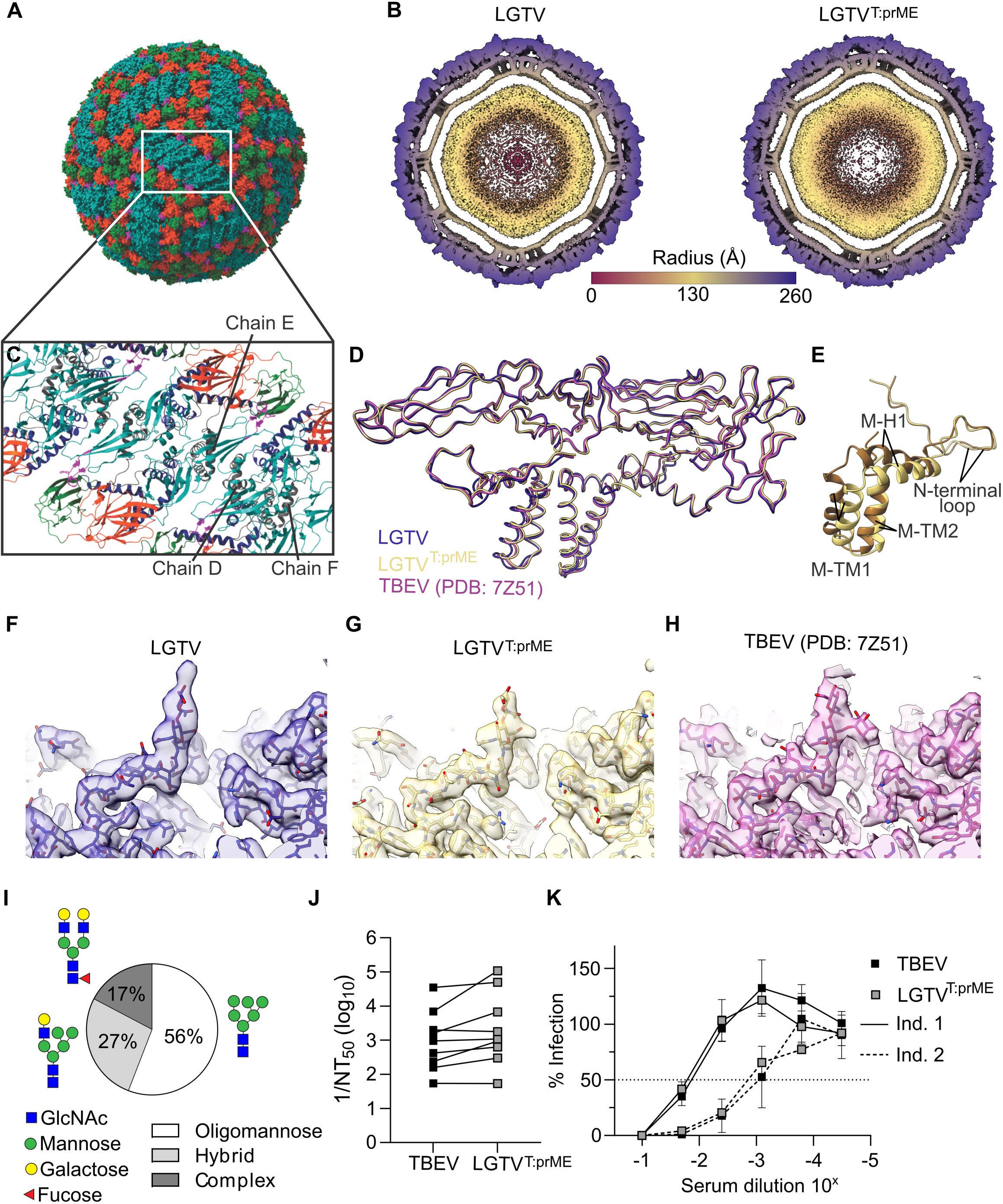
Structural comparison of chimeric LGTV^T:prME^ with LGTV and TBEV Kuutsalo-14. **A)** Surface organization of LGTV^T:prME^ viewed down a two-fold axis of symmetry. The visible E domains are shown in orange (domain I), turquoise (domain II) and green (domain III). **B)** A 30 Å-thick central section of a radially colored isosurface representation of the LGTV and chimeric virus reconstruction, viewed down a two-fold axis of symmetry. Scale bar in Å. **C)** Cartoon representations of the E-M-M-E heterotetramer shown from the top view. The E domains I-III are colored as in A), while domain IV is blue, the E fusion loop is magenta and the M protein is shown in grey. Three chains of the M protein in the asymmetric unit are identified here. **D)** C-alpha backbone overlay of the two models LGTV PDB ID 9FOJ (blue ribbon) and LGTV^T:prME^ PDB ID 9FK0 (yellow ribbon) together with TBEV Kuutsalo-14 [24] (pink ribbon), aligned on chain B of E protein, that has the lowest RMSD value across the three models. **E)** Chain E of M protein in the asymmetric subunit, located between the rafts in LGTV^T:prME^. Conformation consistent with LGTV TP21 (9FOJ) and TBEV Kuutsalo-14 (7Z51) [24] is shown in yellow (deposited as 9FK0). Alternative conformation is shown in brown (deposited as 9H28). Close-up view of Asn154 glycosylation densities of **F**) LGTV, **G)** LGTV^T:prME^ and **H)** TBEV Kuutsalo [24] with the map shown in transparent isosurface and the atomic coordinates in stick representation. **I)** Distribution of N-linked glycan types (complex/hybrid/oligomannose) present at Asn154 of LGVT^T:prME^. GlcNAc=N-acetylglucosamine. An example of the most abundant structure within each group is included as schematic. **J)** Reciprocal 50% neutralizing antibody titers (1/NT50) against TBEV and LGTV^T:prME^ in sera from nine individuals fully vaccinated against TBEV (FMSE-IMMUNE®). **K)** Representative serum neutralization curves for two fully vaccinated individuals with highly neutralizing (solid lines) and medium neutralizing (dotted lines) titers. 50% neutralization indicated by dotted line. Data from two independent experiments performed in duplicates.

We observed densities corresponding to *N*-linked glycosylation at the known E glycosylation site, Asn154, in both LGTV and LGTV^T:prME^ models (Figure 2F-G), consistent to what has previously been shown for WT TBEV [8, 24]. Mass spectrometry analysis of LGTV^T:prME^ confirmed the presence of *N*-linked glycosylation at position Asn154 of the E-protein. When grown in A549 *MAVS^-/-^*cells, LGTV^T:prME^ particles contained different glycan structures at this position, identified as complex-type (17%), hybrid structures (27%) and oligomannose-type 56% (Figure 2H, Supplementary Table 4). The main complex-type glycan was biantennary digalactosylated with core fucosylation. GlcNAc with core fucosylation, whereas the LGTV and LGTV^T:prME^ resolve only the innermost GlcNAc of the *N*-linked glycan at position N154 (Figure 2F-G). Among the complex type glycans, 6% carried one or more sialic acids, but no evidence of fucose linked to the antenna was found. The TBEV strain 93/783 that was used to generate the chimeric virus has an unusual amino acid substitution on the surface of the E protein (Ala83Thr) [9], which is a putative site for *O*-linked glycosylation, however, we did not detect *O*-linked glycosylation of TBEV prM or E ectodomains in viruses produced in A549 *MAVS^-/-^*cells or in primary murine neurons. Furthermore, no corresponding densities were observed in the structural reconstructions.

To confirm that the structural similarity also conferred functional equivalence, we evaluated the neutralizing capacity of serum from nine individuals fully vaccinated against TBEV (FSME-IMMUNE®) against LGTV^T:prME^ and TBEV. We found reciprocal 50% neutralization titers (1/NT_50_) to be similar for TBEV and the chimeric virus (Figure 2I) and observed a similar neutralization profile against the two viruses for both strongly and medium neutralizing serum (Figure 2J). Taken together, these results confirm the structural and antigenic authenticity of the chimeric LGTV^T:prME^.

### Chimeric LGTV^T:prME^ remains non-pathogenic in WT mice and is less pathogenic than LGTV in *Mavs ^-/-^* mice

Next, we investigated the pathogenicity of the chimeric LGTV^T:prME^ *in viv*o. Following infection of C57Bl/6 WT mice with 10^4^ FFU via intraperitoneal (i.p.) injection, neither LGTV^T:prME^ nor LGTV infected mice showed any signs of disease (Figure 3A). In contrast, all mice infected with TBEV succumbed to the infection within 14 days. We have previously shown that *Mavs^-/-^* mice are more susceptible to LGTV infection [25], and may therefore be a suitable model for studying low-pathogenic flaviviruses. These mice reached humane endpoint within seven days p.i for LGTV, independent of the dose (10^4^ and 10^5^ FFU, i.p. injection), while mice infected with LGTV^T:prME^ showed prolonged survival with a median survival of 13 days and nine days after infection with 10^4^ and 10^5^ FFU, respectively (Figure 3B-C). The slight attenuation of the chimeric virus compared to LGTV could be the result of decreased peripheral replication, reduced ability to invade the CNS (neuroinvasion), or reduced neurovirulence. However, we did not observe a difference in neurovirulence between LGTV and LGTV^T:prME^, as measured by survival following intracranial infection of *Mavs^-/-^* mice (Figure 3D).

**Figure 3.**
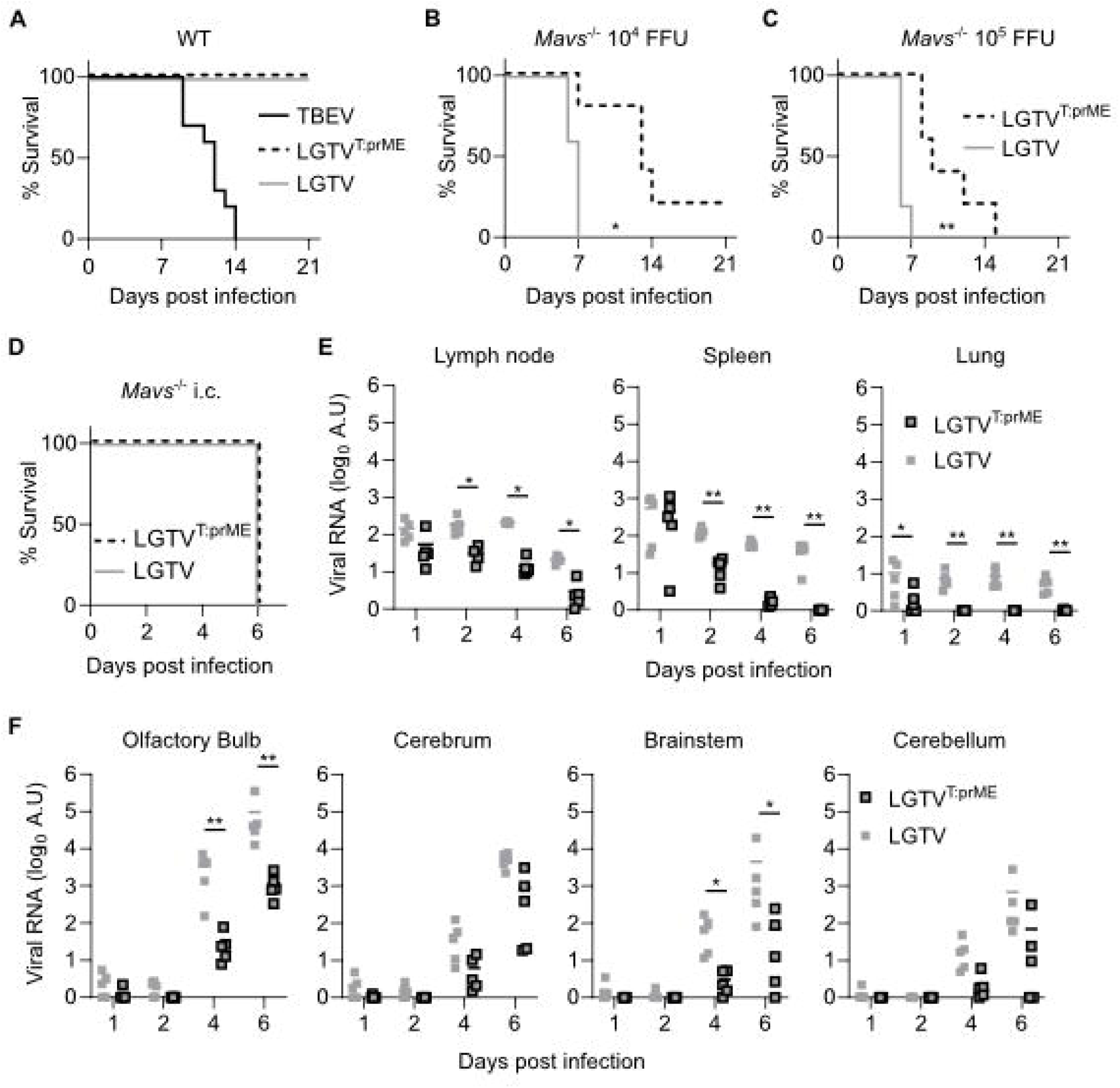
*In vivo* characterization of chimeric LGTV^T:prME^. **A)** Survival analysis of C57Bl/6 WT mice infected intraperitoneally (i.p.) with 10^4^ FFU of TBEV (n=10), LGTV^T:prME^ (n=4) or LGTV (n=5) and monitored for up to 21 days. Survival analysis of *Mavs^-/-^* mice infected by i.p. injection with **B)** 10^4^ FFU or **C)** 10^5^ FFU of the indicated virus (n=5 per group). **D)** Survival analysis of *Mavs^-/-^*mice infected intracranially (i.c.) with 10^2^ FFU of LGTV^T:prME^ or LGTV (n=5 per group). Survival differences were tested for statistical significance by the log-rank test, * pL<L0.05, ** p < 0.01. *Mavs^-/-^* mice infected i.p. with 10^5^ FFU of LGTV^T:prME^ or LGTV and viral loads quantified by RT-qPCR in **E)** peripheral organs; lymph nodes, spleen and lung or **F)** different regions of the brain; olfactory bulb, cerebrum, brainstem and cerebellum. Statistical significance calculated using Mann-Whitney test, * p < 0.05, **p < 0.01.

To better understand the observed difference in pathogenicity following i.p. infection, we assessed viral loads in various organs on days 1, 2, 4, and 6 p.i with 10^5^ FFU chimeric LGTV^T:prME^ or LGTV. Peripheral organs (lymph nodes, spleen, lung) showed viral RNA from day 1, with levels remaining stable or slightly decreasing (Figure 3E). By day 2, the chimeric virus showed significantly lower viral RNA load than LGTV (Figure 3E). Viral levels in the brain increased on day 4, and then further to day 6, with the chimeric virus showing significantly lower levels in the olfactory bulb and brainstem at both time points (Figure 3F). From these data, it is not possible to exclude a difference in neuroinvasion between LGTV^T:prME^ and LGTV, but it is more likely that the differences observed in peripheral replication are responsible for the lower viral loads detected in the brain of animals infected with the chimeric virus.

### LGTV and chimeric LGTV^T:prME^ predominantly infect cerebral cortex while TBEV highly infects cerebellum

Genetic variance in the E protein of TBEV impacts neurovirulence [9], but its role in determining tropism is less known. Using the neurovirulent chimeric virus, we evaluated the structural proteins’ role in whole brain distribution and tropism. Therefore, brains were collected at humane endpoint (5-6 days p.i., Supplementary Figure 2A) and analyzed using 3D whole brain OPT imaging. In general, TBEV exhibited significantly higher overall infection rates in the brain compared to LGTV and LGTV^T:prME^, as observed in both maximum intensity projection (MIP) images and whole-brain quantification (Figure 4A-B, Supplementary Figure 2B, Supplementary Video 1-3). TBEV predominantly infected the cerebellum, whereas the majority of LGTV and LGTV^T:prME^ infection was localized in the cerebrum.

**Figure 4.**
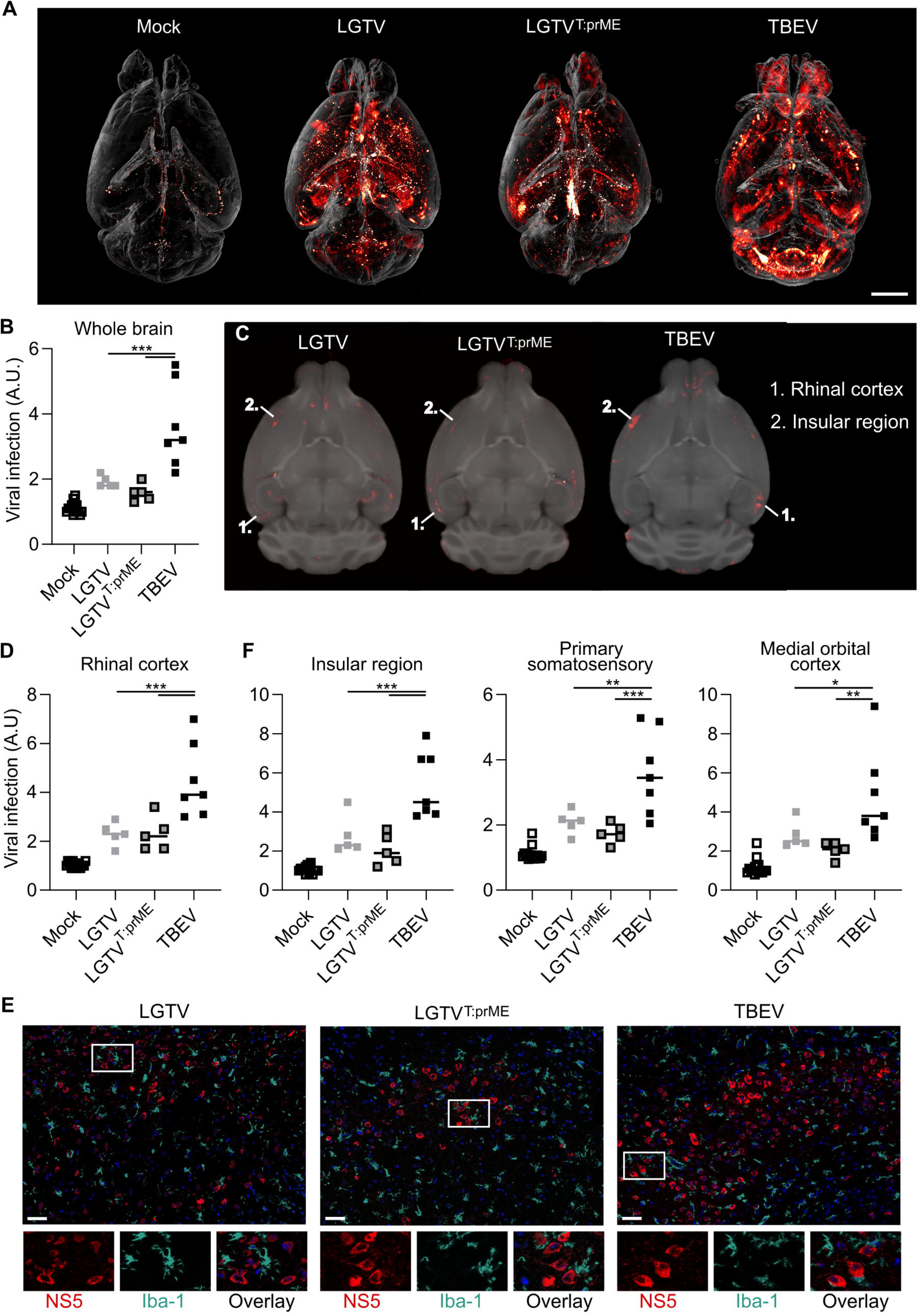
Distribution and tropism of chimeric LGTV^T:prME^ compared to parental viruses in mouse brain. **A)** Virus infection in whole mouse brain visualized by OPT. Maximum intensity projection (MIP) from mice infected by i.c. inoculation with LGTV (10^4^ FFU, n=5), LGTV^T:prME^ (10^4^ FFU, n=5) or TBEV (10^2^ FFU, n=7), stained against viral NS5 protein. Scale bar 2 mm. Quantification of viral OPT signal in **B)** whole brain, or **D)** and **F)** selected or composite regions of interest (ROIs). Statistical significance calculated using ordinary one-way ANOVA with Tukey’s multiple comparison test, ** p < 0.01, *** p < 0.001. **C)** Anatomical mapping (axial plane) of viral infection from OPT signal (red) co-registered to the OCUM brain template (grey). E) Axial brain sections stained against viral NS5 (red) and Iba-1 (turquoise), nuclei stained with DAPI are blue. Images taken by confocal microscopy, magnification 40x. Scale bar 50 μm.

To study the distribution of these viruses with more anatomical detail, we created fusion images displaying viral signal in the anatomical context of an MR-based brain template (Supplementary Video 4-6 (axial plane). These whole brain OPT-MR fusion images allowed us to map and quantify viral brain distribution in all 336 anatomical regions of the OCUM brain atlas (Supplementary Data 1) [26]. For LGTV and the chimeric virus, the brain regions with the most infection were localized in the cerebral cortex with most pronounced infection rates in the rhinal cortex (comprising of the entorhinal, perirhinal and ectorhinal cortices) (Figure 4C-D, Supplementary Table 5, Supplementary Video 7). This infection pattern was also confirmed by confocal microscopy, where most infected cells displayed neuron-like morphology (Figure 4E). Our confocal imaging data indicate that microglia are highly activated following infection, as evidenced by their morphology and number of Iba-1 positive cells as compared to mock infected mice (Figure 4E, Supplementary Figure 3A). Microglia are often observed in close proximity to infected cells but remain resistant to the infection by all three viruses (Figure 4E). Furthermore, both viruses displayed infection in the somatosensory cortex, insular region and medial orbital cortex (Figure 4D, Supplementary Video 8). Interestingly, these regions were also among the most highly infected cerebral regions in TBEV infected brains (Supplementary Table 5), indicating similar distribution of all three viruses in cerebral cortex. The only white matter tract among the highest infected brain regions of LGTV and LGTV^T:prME^ was the olfactory limb of the anterior commissure (Supplementary Table 5).

Apart from cerebellar vermis lobules 1-2 (lingula and ventral central), cerebellar infection was low and not significant for LGTV and the chimeric virus, which contrasts strikingly with TBEV, where we observed infection in most cerebellar regions (Figure 5A, Supplementary Table 5, Supplementary Video 9). The OPT-template fusion images reveal a distinct pattern of TBEV infection in the cerebellum (Figure 5B), with the highest viral signal in the cerebellar grey matter (Figure 5C). Since OPT lacks the resolution to visualize the distinct layers of the cerebellar grey matter, we used confocal microscopy for further characterization. We observed TBEV infection in the Purkinje, molecular and granular cell layers of the cerebellar grey matter (Figure 5D). TBEV-infected cells predominantly overlapped with Calbindin C staining, clearly indicating TBEV’s preference for infecting Purkinje cells in the cerebellum. Given the limited number of cells displaying viral antigen in LGTV^T:prME^ infected brains, this preference cannot solely be attributed to the structural proteins (Supplementary Figure 3B). Other regions that were highly infected with TBEV but not with the two other viruses include the pons, medulla and hypothalamus (Supplementary Figure 3C, Supplementary Table 5). Taken together, the distribution patterns of LGTV and the chimeric virus are strikingly similar, whereas TBEV shows a stronger and more widespread infection, especially in the cerebellum.

**Figure 5.**
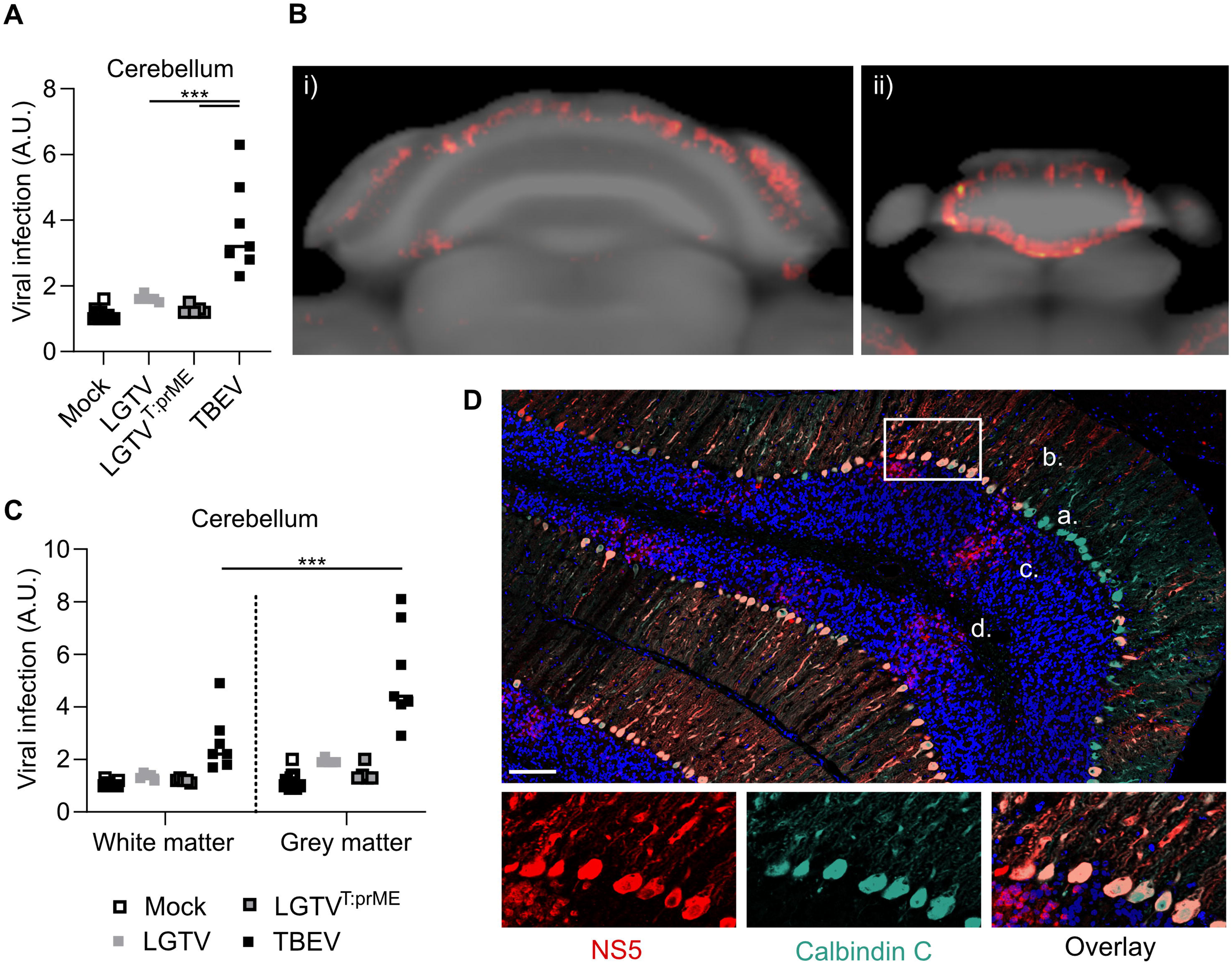
High infection of Purkinje cells in the cerebellar grey matter by TBEV. **A)** Quantification of viral OPT signal in cerebellum of mice infected by i.c. inoculation with LGTV (10^4^ FFU, n=5), LGTV^T:prME^ (10^4^ FFU, n=5) or TBEV (10^2^ FFU, n=7), stained against viral NS5 protein. **B)** Axial view of cerebellum of an TBEV infected mouse, showing viral infection from OPT signal (red) co-registered to the OCUM MRI based brain template (grey), i) cerebellum, ii) cerebellar vermis. **C)** Quantification of viral OPT signal in composite region of interests (ROIs) in white matter and gray matter of the cerebellum. Statistical significance calculated using ordinary one**-**way ANOVA with Tukey’s multiple comparison test, *** < 0.001. **D)** Axial brain sections stained against viral NS5 (red) and Calbindin C (turquoise), nuclei stained with DAPI are blue, a. Purkinje cell body, b. molecular layer, c. granular cell layer, d. white matter. Images taken by confocal microscopy, magnification 40x. Scale bar 100 μm.

## Discussion

Here, we characterized and utilized a chimeric virus containing the prM and ecto-E proteins from TBEV within the genetic background of low-pathogenic LGTV. This allowed us to specifically study the role of TBEV structural proteins in determining the distribution and tropism of viral infection in the brain. Importantly, in line with similar previous studies, the chimeric virus remained low-pathogenetic, as shown by non-lethality in adult C57Bl/6 WT mice and prolonged survival compared to LGTV in immunocompromised (Mavs^-/-^) mice. Comparison of viral distribution in the brain across the three viruses shows very similar patterns in cerebral cortex, with the rhinal cortex being the primary site of cortical infection for all viruses. This indicates that LGTV and TBEV either use the same entry receptor(s) or that their receptors have overlapping cell distributions. In general, TBEV showed more widespread and higher infection compared to the other viruses and a particularly strong propensity to replicate in the cerebellum. This shows that the structural proteins alone do not determine viral tropism and, suggesting that the non-structural proteins may play a role in shaping tropism, likely through their immunomodulating role.

Analysis of virus growth kinetics *in vitro* showed that viral replication, progeny virus production and cytopathic effects are mainly determined by the non-structural proteins. Meanwhile, the chimeric virus showed an advantage over parental LGTV in cell-to-cell spread, suggesting a difference between TBEV and LGTV prM-E proteins in this aspect. Similarly, a study comparing TBEV strains with different pathogenicity showed a role of the structural proteins in non-viremic tick-to-tick transmission, while the virus’s ability to induce cytopathic effect *in vitro* relied mainly on the non-structural proteins [27]. In line with this, we observed a clear cytopathic effect for TBEV 72 h p.i, but not for the chimeric virus.

Chimeric TBEVs have previously been developed by inserting TBEV prM and E proteins into various flavivirus backgrounds, such as YFV 17D, DENV2/4, WNV, JEV SA14-14-2, and LGTV [19–21]. Earlier work on TBEV/LGTV chimeras has primarily focused on vaccine development and reducing neuropathology [22, 28], but provided limited characterization of the generated viruses beyond basic lethality. We confirm here that the chimeric LGTV^T:prME^ remained non-pathogenic in adult C57Bl/6 WT mice. In immunocompromised (*Mavs^-/-^*) mice, we observed similar disease progression following i.c. infection with LGTV and chimeric LGTV^T:prME^, but prolonged survival of the chimeric virus infected animals after i.p. infection. Similarly, data from 3-week-old mice, infected with a chimeric TBEV/LGTV, show LD_50_ values comparable to LGTV following i.c. infection, but higher LD_50_ after peripheral infection with the chimeric virus [22]. We analyzed viral loads in various peripheral organs and brain regions following i.p. infection, and could observe lower viral loads for the chimeric virus in all analyzed peripheral organs already at day 2 p.i. This suggests that the attenuation of LGTV^T:prME^ occurs already during peripheral replication or through more efficient viral clearance by the early immune response. Infection of cultured murine macrophages by two TBEV strains with different pathogenicity showed strain-specific differences in infection and immune activation [15], but the mechanism and viral proteins involved remain to be further characterized. The observed attenuation might also be due to sub-optimal interactions between the prM protein from TBEV and the C protein or transmembrane domain of the E protein of LGTV.

We and others have previously shown that genetic variation in the E protein affects antibody binding and neutralization [8, 9, 29, 30], which could influence the efficiency of available vaccines. Genetically divergent TBEV strains (>10% nucleotide sequence difference) and overlapping endemic areas for related tick-borne flaviviruses (TBFVs) highlight the need to understand cross-reactivity. The structural similarity and coherence of neutralizing titers against LGTV^T:prME^ and TBEV suggest that low-pathogenic chimeric viruses could facilitate this research. We also confirmed the presence of an *N*-linked glycan at the previously known glycosylation site N154 on the E protein, which is known to influence virus secretion and pathogenicity in mice [31]. The heterogeneous glycan composition at this site comprises a substantial degree of oligomannose and biantennary complex type glycans carrying core fucosylation, in line with previously published data for WT TBEV E protein [32].

From the present data, it is clear that the infection pattern of the three viruses is very similar in cerebrum and more specifically, in cerebral cortex. The most apparent difference between TBEV and LGTV^T:prME^ is the substantial cerebellar infection in TBEV-infected mice, while mice infected with the chimeric virus showed little to no cerebellar infection. Cerebellar infection was not apparent in WT or immunosuppressed *Ifnar*^-/-^ mice infected with LGTV [18, 33]. We can only speculate on the reasons for this, but one possibility might be the rapid disease progression in *Ifnar*^-/-^ mice, which reduces the time for dissemination. Looking closer at the cerebellum, confocal microscopy revealed extensive TBEV infection in Purkinje cells. These are large inhibitory neurons found in cerebellar grey matter and regulate motor function. Interestingly, TBEV infection of Purkinje cells has earlier been detected in human post-mortem autopsy samples [11]. Infection of these cells by TBEV, but not the chimeric virus, indicates a mechanism other than receptor specificity and warrants further investigation. Additionally, whether or not Purkinje cells become infected following other routes of inoculation, which include viral entry into the brain from the periphery, remains to be discovered. We also observed infection in the medulla and pons with TBEV, regions which have been identified to be infected in post-mortem brain tissue from patients with TBE [11]. Interestingly, the pons was also infected in LGTV, though they did not rank among the top 20 most infected regions. In contrast to our study, analysis of post-mortem brain tissue from TBE patients observed only limited infection in the cerebral cortex. However, this could also be due to their focus on a specific part of the much larger cortex found in humans. Our whole brain imaging approach with atlas coregistration provides more detailed information, as it includes and distinguishes between the many individual regions of the cerebral cortex.

In conclusion, we have developed a low-pathogenic model virus to study the structural proteins E and prM of TBEV. Using this model, we demonstrate that the distribution and tropism of LGTV and TBEV in the brain are not solely dependent on receptor tropism. The primary difference observed with TBEV, compared to the other viruses, is its strong propensity to infect the cerebellum and especially Purkinje cells. Together, this deepens our understanding of the structural proteins’ role as pathogenicity determinants of tick-borne flaviviruses.

## Materials and methods

### Cells and reagents

BHK-21 (ATCC CCL-10) and A549 (ATCC-CCL-185) cells were cultured in Dulbecco’s modified Eagle’s medium (DMEM, HyClone^TM^, Logan, UT, USA) with 5% fetal bovine serum (FBS, HyClone^TM^), 20 units/ml penicillin, and 20 μg/ml streptomycin (PeST, Gibco™, Billings, MT, USA) at 37°C in 5% CO_2_. A549 cells with a genetic knockout of mitochondrial antiviral signaling protein (MAVS), annotated A549 *MAVS^-/-^* were a kind gift from Gisa Gerhold [34] and were cultured in DMEM with 10% FBS, penicillin and streptomycin as described above and 2 µg/ml puromycin (Gibco™).

### Generation of TBEV and LGTV infectious clones and design of chimeric virus

Infectious cDNA (icDNA) clones of LGTV strain TP21 (LGTV) (NCBI Reference Sequence: NC_003690.1) and TBEV strain 93/783 (TBEV) [23] (GenBank Accession no MT581212.1) were constructed in pCC1BAC^TM^ vector (EPICENTER). Briefly, cDNAs of LGTV and TBEV were placed between sequences of promoter SP6 RNA polymerase at the 5’ end of the construct and negative-strand ribozyme of hepatitis delta virus at the 3’ end. The sequence of each DNA was split into 5 fragments flanked by naturally occurring unique restriction sites; fragments were obtained as synthetic DNAs (Genscript, USA) and assembled into full-length icDNAs using four step ligation protocols. All plasmids containing intermediate cloning products and icDNAs were propagated using EPI300TM *E. coli* cells (LGC, Biosearch Technologies, USA).

A chimeric clone containing the nucleotide (nt) sequence coding for prM and ecto-E of TBEV strain 93/783 in a LGTV background (LGTV^T:prME^) was generated based on the icDNA clone of LGTV. The fragment between restriction sites MluI and AatII (LAN1) corresponding to the region encoding TBEV prM and the ectodomain of E (nt 505-2289) was synthesized and cloned into a pUC57Kan vector backbone by service provider (Genscript). Chimeric LAN1 fragment was used to replace the corresponding fragment of pCCI-LGTV using standard molecular cloning procedures; sequences of obtained clones were verified by Sanger sequencing.

### Rescue and propagation of recombinant viruses

Plasmid containing icDNAs were linearized with SmiI FastDigest enzymes (Thermo Fisher Scientific, Waltham, MA USA) followed by DNA purification using DNA Clean & Concentrator-5 kit (BioSite). 0.05-0.1 µg of linearized DNA were used as input in a 5 µl mMESSAGE mMACHINE SP6 in vitro transcription reaction (Invitrogen). BHK-21 cells grown in 6 well cell culture plates (2×10^5^ cells/well) were transfected with in vitro-transcribed RNAs using Lipofectamine 2000 reagent (Invitrogen). Supernatant from the transfected cells (passage 0) was collected over time and titrated on A549 *MAVS^-/-^* cells by focus forming assay, as described below. The earliest collected sample showing a high titer (>10^5^ focus forming units (FFU)/ml) was used to generate a virus stock by infection of A549 *MAVS^-/-^* cells at multiplicity of infection (MOI) 0.005. These stocks (passage 1) were sequenced and used for all downstream experiments without further passaging.

### Virus infections and focus forming assay

A549 cells grown in 24 well cell culture plates (1.5×10^5^ cells/well) were infected at indicated MOI for 1 h before the inoculum was replaced with fresh media (DMEM, PeSt and 2% FBS) and cells were incubated at 37°C in 5% CO_2_. Viral replication was quantified by reverse transcription quantitative PCR (RT-qPCR) and the release of infectious virus was quantified by focus forming assay.

For quantification of infected cells, monolayers of A549 cells, grown in 96 well cell culture plates (10,000 cells/well) were infected with the indicated virus at a MOI 0.01. Following 1 h inoculation, the media was replaced by DMEM containing PeSt and 2% FBS and cells were incubated for an additional 48 h or 72 h. Plates were washed once with PBS, fixed with 4% formaldehyde for 30 min and permeabilized for 15 min at room temperature with PBS containing 20 mM glycine and 0.5% Triton-X100. Plates were stained with primary antibodies targeting E-protein (mouse monoclonal antibody 1786 [35], dilution 1:1000) and NS5 protein (chicken polyclonal IgY, in-house [18], dilution 1:500) for 1 h at room temperature followed by treatment with secondary antibodies donkey anti-mouse Alexa Fluor™ 647 and goat anti-chicken IgY Alexa Fluor™ 555 (Invitrogen, AB_162542 and AB_2535858) in PBS containing 10% FBS, 0.05% Tween-80 and 0.1 µg/ml DAPI. Images were obtained on a Cytation^TM^ 5 (BioTek, Winooski, VT, USA) and the percentage of infected cells was calculated as double-positive cells divided by cell count by DAPI.

For focus forming assay, monolayers of A549 *MAVS^-/-^*cells, grown in 96 well cell culture plates (25,000 cells/well) were infected with 1:10 dilution series of the indicated virus. Following 1 h inoculation, the media was replaced by a semisolid overlay (DMEM, PeSt, 2% FBS and 1.2% Avicel RC-581 (FMC Biopolymer)) and cells were incubated for an additional 48 h or 72 h. Plates were washed, stained and imaged as described for quantification of infected cells. Plaques were visualized with primary antibodies targeting the E-protein followed by treatment with donkey anti-mouse Alexa Fluor™ 647 as a secondary antibody.

### Virus passaging and sequencing

Serial passaging was performed for LGTV and LGTV^T:prME^ using A549 cells grown in 12 well cell culture plates (200,000 cells/well). Cells were infected at MOI 0.1 for 1h before the inoculum was replaced with fresh media (DMEM, PeSt and 2% FBS). After 72h, the supernatant was collected, titrated and subsequently used as inoculum for the next step. For each virus, this process was repeated 10 times in triplicates. 100 μl of the supernatant from the last round (p10) or the original virus stock (p1), was used for extraction of viral RNA using the QIAmp viral RNA kit (Qiagen) and 10 μl of the eluate was used as input for cDNA synthesis using the high-capacity cDNA reverse transcription kit (Applied Biosystems). The cDNA was diluted 1:10 and 1 μl used as input for the PCRs using KOD Hot Start DNA Polymerase (MilliporeSigma), 50 μl reactions, 35 cycles, 52°C annealing temperature. A total of seven amplicons spanning the entire coding region were designed for LGTV and adjusted for LGTV^T:prME^. Amplicons were gel purified (Gel extraction kits E.Z.N.A., Omega Bio-Tek) and pooled in equimolar amounts. The sequencing libraries were prepared using the rapid barcode kit SQK-RBK110-96 (Oxford Nanopore, UK) according to the manufacturer’s instructions. Concentrations of the resulting barcoded libraries were measured using Qubit 1× dsDNA high-sensitivity assay kit (Invitrogen) and pooled in equimolar amounts before sequencing on a R9.4.1 FLO-MIN106D flow cell. Using MinKNOW (v. 22.08.4), the sequencing ran for 15 h until a sufficient yield had been reached.

The raw fast5 signal data files generated by MinKNOW were base called and demultiplexed using guppy v. 6.4.8 and the sup basecalling model. A custom bioinformatic pipeline utilizing the Snakemake (v. 7.19.1) workflow management system was used to analyze the demultiplexed FASTQ files for each sample. In brief, the pipeline consisted of analysis of quality and size distribution of the reads using NanoPlot (v. 1.41.6). Adapter trimming was performed using Porechop (v. 0.2.4), and de novo assembly of the adapter-trimmed reads was conducted using Flye (v. 2.9.2). Due to the short length of the reads, Flye did not manage to create contigs spanning the whole genome of the viruses. The partial contigs were therefore stitched together using the scaffold command in Ragtag (v 2.1.0). Next, the reads were mapped to the appropriate reference genome for each sample using Minimap2 (v. 2.26). The resulting BAM file was used as input for SAMtools (v. 1.17) mpileup, and a consensus sequence was generated using iVar (v. 1.4.2). Coverage plots for each sample were produced using bam2plot (https://github.com/willros/bam2plot). To inspect the per-base coverage and identify interesting regions of the BAM file, the program perbase (v. 0.9.0) was used alongside in-house scripts.

### Virus infections of mice

Six to 12-week-old WT C57BL/6 mice and MAVS knockout mice (*Mavs^-/-^*) of mixed gender in C57BL/6 background (kind gift of Nelson O Gekara, Umeå University) were used for the infection experiments. For intraperitoneal (i.p.) infections, mice were injected with indicated viral dose diluted in 100-200 µl PBS, either following brief sedation with diethyl ether (TBEV) or without prior sedation (LGTV, LGTV^T:prME^). For intracranial (i.c) infections, mice were sedated with 4% isoflurane or by intramuscular ketamine-xylazine anesthesia before inoculation of indicated dose diluted in 20 μl of PBS or PBS only (mock). Mice were euthanized using CO_2_ or sedated followed by cardiac perfusion at the previously described criteria for humane endpoint [18].

### RNA isolation and RT-qPCR

Total RNA was extracted from cells or organs and the amount of viral RNA was quantified by RT-qPCR as previously described [25]. Briefly, for organs, mice were perfused with 20 ml of ice cold PBS and the collected organs were homogenized in QIAzol Lysis Reagent (Qiagen) using 1.3 mm Chrome-Steel Beads (BioSpec) and the FastPrep-24 homogenizer (MP). RNA was obtained by phenol-chloroform extraction followed by purification using Nucleo-Spin RNA II kit (Macherey-Nagel) according to the manufacturer’s instructions. For cell cultures, RNA was extracted using Nucleo-Spin RNA plus kit (Macherey-Nagel). High-capacity cDNA Reverse Transcription kit (Thermo Fisher) was used for cDNA synthesis with 500-1 000 ng of total RNA as input. Viral RNA was quantified with qPCRBIO probe mix Hi-ROX (PCR Biosystems) and primers recognizing LGTV NS3 gene [25] or TBEV 3’ UTR [36]. Mouse GAPDH or human actin mRNAs were used as a reference and detected by QuantiTect primer assay (GAPDH; QT01658692, Actin; QT01680476, Qiagen) and the qPCRBIO SyGreen mix Hi-ROX (PCR Biosystems). All experiments were run in technical duplicates on a StepOnePlus real-time PCR system (Applied Biosystems).

### Sample preparation cryo-EM

A549 *MAVS^-/^*^-^ cells at 95% confluency in T175 flasks were infected by LGTV^T:prME^ or LGTV at MOI 0.1 as follows: the cells were washed 4 times with PBS, virus was added in infection medium (VP-SFM (Gibco, Waltham, MA, USA) + 2 mM glutamine + 35 nM rapamycin), and incubated at 37°C in a 5% CO_2_ atmosphere. At 72 h p.i, the supernatant was collected and centrifuged for 5 min at 4°C at 3100× g followed by high-speed centrifugation for 1 h at 13200 x *g* at 4°C. The supernatant was collected, 4 µL of Benzonase^®^ Nuclease (Merck) was added, and the mixture was incubated for 10 min at room temperature. After benzonase treatment, the supernatant was added to a suspension of Capto™ Core 700 beads (medium:beads of 4:1) in HNE buffer (20 mM HEPES + 150 mM NaCl + 1 mM EDTA, pH 8.5), and was incubated on a rotary shaker at 4°C for 30 min and cleared at 4°C for 5 min at 120 x g. The Capto™ Core 700 purification was repeated once. For the LGTV, the supernatant after Capto™ Core 700 purification was filtered through a Corning™ Disposable Vacuum Filter system (0.22 µm). The supernatant was concentrated using Amicon Ultra-15 centrifugal filter units (MWCO 100kDa) for 20 min at 4°C at 4500 x *g*. The concentrated sample was inactivated by irradiation with 0.1 J/cm2 UV-C using a UVP Crosslinker (Analytik, Jena, Germany). Inactivated LGTV^T:prME^ was vitrified without further preparation. LGTV was additionally concentrated by pelleting in a Beckmann Airfuge^®^ for 8 min at 29 psi in A-100/18 rotor at room temperature. The pellet was resuspended in 13 µL of inactivated sample after ultrafiltration. Samples were vitrified on glow-discharged QUANTIFOIL® R 1.2/1.3, 2 nm continuous carbon on 300 mesh Cu grids, using a Leica EM GP plunger at 80% humidity and 1.5 s blotting time using front blotting. For LGTV, the sample was incubated on the grid for 30 seconds prior to plunging.

#### Cryo-EM data collection, reconstruction and model building

The data for LGTV were collected at the University of Helsinki using an FEI Talos Arctica microscope equipped with a Falcon 3 detector. The microscope was operated in the counting mode at 200 kV. A total of 1430 movies were collected at a nominal magnification of 150,000x at a 0.96 Å^2^/pixel sampling rate with an acquisition area of 4096 by 4096 pixels and a total dose of 40 electrons/Å^2^ divided over 40 frames. The data for LGTV^T:prME^ were collected at the SciLifeLab facilities in Umeå, Sweden, using an FEI Titan Krios microscope equipped with a Falcon 4i detector. The microscope was operated in the counting mode. A total of 32,122 images were collected at a nominal magnification of 130,000× at a 0.917 Å/pixel sampling rate with an acquisition area of 4096 by 4096 pixels and a total dose of 40 electrons/Å^2^. Movies were recorded in the EER format.

Image processing was performed at the CSC-IT Center for Science Ltd. and in-house, using the CryoSPARC framework [37]. The micrographs were aligned using CryoSPARC’s patch motion correction function, and their respective CTF functions were estimated with patch CTF estimation [37]. Particles were picked using the template picker function, using the template generated from classified blob picks [37]. The initial model was generated using Cryosparc’s ab-initio reconstruction, using C1 symmetry and then refined, using the Non-Uniform Refinement protocol in CryoSPARC [39], with the imposition of I2 symmetry. The maps were postprocessed using DeepEMhancer [40]. Initial atomic models were generated as M3E3 complexes using AlphaFold plugin in PHENIX [41, 42]. The resultant models were flexibly fitted into the sharpened maps using ISOLDE integrated into ChimeraX [43]. The model was real-space refined using Phenix [44], after which clashes were fixed in ISOLDE with model-based distance and torsions constraints imposed [45]. Root-mean-square deviations (RMSD) between the refined models and TBEV stain Kuutsalo-14 (PDB ID 7Z51, [24]) were calculated in ChimeraX [43].

### Glycosylation analysis by mass spectrometry

A549 *MAVS^-/-^* cells grown in DMEM containing PeSt and 0.35 µM Rapamycin (Selleck Chemicals) without serum were infected with LGTV^T:prME^ at MOI 0.01. At day 4 p.i supernatant was harvested, floating cells were removed by centrifugation at 500 x g for 5 min, after which virus was collected by ultracentrifugation using an Optima L-80 XP ultracentrifuge (Beckman Coulter) at 100,000 x g in a Sw 41 Ti rotor. Purified viral particles (estimated amount of 10 µg) were resuspended in 500 µl PBS supplemented with 2% sodium dodecyl sulfate (SDS, Merck), pH 7.4, and reduced with 100 mM dithiothreitol (DTT, Thermo Fisher Scientific) for 30 min at 56°C followed by processing using a modified filter-aided sample preparation (FASP) method [43]. In short, the reduced samples were diluted to 1:4 in 8 M urea solution, transferred onto a Pall Nanosep centrifugation filter (Pall Corporation), washed repeatedly with 8 M urea and once with digestion buffer (0.5% sodium deoxycholate (SDOC, Sigma-Aldrich) in 50 mM triethylammonium bicarbonate (TEAB, Fluka)). Alkylation of the samples was performed with 18 mM 2-iodoacetamide (IAM) for 30 min at room temperature in the dark, followed by an additional wash with digestion buffer.

Samples were digested with trypsin (0.3 µg, Pierce MS grade Trypsin, Thermo Fisher Scientific) at 37°C overnight, an additional portion of trypsin (0.3 µg) was added followed by an additional 3 h incubation. Proteolytic digestion was performed in digestion buffer using chymotrypsin (first incubation overnight at 23 °C and second incubation for 3 h at 23°C, both with 0.2 µg chymotrypsin). Proteolytic peptides were collected by centrifugation and the filter was additionally washed with 35 µl of 50 mM TEAB to ensure that all material was eluted from the filter. Collected peptides were purified using HiPPR™ spin column, followed by acidification with 10% trifluoroacetic acid to remove the remaining SDOC. The collected supernatant was desalted using Pierce™ peptide desalting spin columns prior to LC-MS/MS analysis. All sample purification procedures were performed according to the manufacturers’ instructions. The detergent-free preparation was dried down and reconstituted in 20 µl of 2% acetonitrile (ACN, Merck), 0.1% formic acid (FA, WVR) for LC-MS analysis.

The sample was analyzed on an Orbitrap Exploris 480 mass spectrometer interfaced with Easy-nLC1200 liquid chromatography system (both Thermo Fisher Scientific). Peptides were trapped on an Acclaim Pepmap 100 C18 trap column (100 μm x 2 cm, particle size 5 μm, Thermo Fisher Scientific) and separated on an in-house packed analytical column (75 μm x 30 cm, particle size 3 μm, Reprosil-Pur C18, Dr. Maisch) using a gradient from 5% to 35% ACN in 0.2% formic acid over 75 min at a flow of 300 nL/min. Each preparation was analyzed using two different MS1 scan settings, m/z 380-1500 and m/z 600-2000, both at a resolution of 120K. MS2 analysis was performed in a data-dependent mode at a resolution of 30K, using a cycle time of 2 seconds. The most abundant precursors with charges 2-7 were selected for fragmentation using high energy collision dissociation at collision energy settings of 30. The isolation window was set to either 1.2 m/z for data acquired in m/z 380-1500 range or 3.0 m/z for data acquired in m/z 600-2000 range. The dynamic exclusion was set to 10 ppm for 20 s.

The acquired data were analyzed using Proteome Discoverer 2.4 (Thermo Fisher Scientific). Database searches were performed against custom protein database, consisting of selected TBEV protein sequences and SwissProt Human database to control for the production cell-line protein contamination. The data acquired in m/z 380-1500 range were searched using Sequest HT. Precursor mass tolerance was set to 10 ppm and fragment mass tolerance to 20 mmu. Chymotryptic peptides with up to five missed cleavages were accepted. Methionine oxidation and Ser/Thr phosphorylation were set as variable modifications. Cysteine alkylation was set as fixed modifications. Target decoy was used for peptide spectrum match (PSM) validation.

For glycopeptide analysis, data were searched using Byonic search engine (Protein Metrics, San Carlos, CA). Precursor mass tolerance was set to 10 ppm and fragment mass tolerance to 30 ppm. Chymotryptic peptides with up to five missed cleavages were accepted with variable modification of methionine oxidation and fixed cysteine alkylation. Custom O-glycan database consisting of 9 compositions was used to search for O-glycosylated peptides in the dataset acquired at m/z 380-1500. Custom N-glycan database consisting of 129 compositions was used to search for N-glycosylated peptides in the dataset acquired at m/z 600-2000. Target decoy was used for PSM validation. Prior to the final assignment, the identified glycosylated peptides were manually validated based on the observed fragmentation pattern, the number of glycoforms per site, the number of PSM per glycoform, and the retention time windows for the different glycoforms (at the same site). The extracted ion chromatogram peak intensities were used to determine the glycoform abundances and are expressed as percent of total signal for all modified and non-modified peptides sharing the same amino acid sequence.

### Neutralization assay

A total of nine anonymized serum samples from the German National Consultant Laboratory for TBEV were included [9]. Sera were from individuals with a complete basic immunization with FSME-IMMUN®, with the last vaccine shot more than 3 months before blood sampling and had never suffered from tick-borne encephalitis (TBE), according to anamnestic information. For the neutralization test, sera were diluted 1:5 in DMEM containing PeSt and complement was inactivated at 56°C for 30 min. Five-fold serial dilutions of serum in DMEM containing PeSt were mixed 1:2 with 150 FFU of TBEV or LGTV^T:prME^ and incubated at 37°C for 30 min before addition to A549 *MAVS^-/-^*cells, grown in 96 well cell culture plates (25,000 cells/well). Following 1 h inoculation, the media was replaced by a semisolid overlay (DMEM containing PeSt, 2% FBS and 1.2% Avicel RC-581 (FMC Biopolymer)) and incubated for an additional 48 h (TBEV) or 72 h (LGTV^T:prME^). Viral foci were revealed by focusing-forming assay and percent inhibition was calculated compared to untreated virus control. The reciprocal 50% neutralization titer (1/NT_50_) for each serum was calculated with a non-linear fit of normalized data using GraphPad Prism 9.

### Whole-mount immunohistochemistry and Optical projection tomography (OPT)

Whole brains from infected or mock-treated mice were isolated, stained and imaged as previously described [18], with minor changes. In brief, following cardiac perfusion with 20 ml of cold PBS and 20 ml of cold 4% paraformaldehyde in PBS, brains were fixed overnight in 4% formaldehyde at 4°C. Brains were dehydrated stepwise with MeOH, permeabilized by repeated freeze-thaw cycles at −80°C and bleached in H_2_O_2_:DMSO:MeOH (3:1:2) for 48 h. Brains were rehydrated into TBS-T (50 mM Tris-HCl, pH 7.4, 150 mM NaCl, and 0.1% v/v TritonX-100), blocked for 48 h at 37°C (10% goat serum, 5% DMSO and 0.01% NaAz in TBS-T), stained with primary (chicken anti-NS5 diluted 1:2000 (TBEV) or 1:1000 (LGTV and LGTV^T:prME^)) and secondary (goat anti-chicken Alexa Fluor 680; diluted 1:1000) antibodies diluted in blocking buffer for 5 days each. The brains were mounted in SeaPlaque™ agarose (#50101, Lonza, Switzerland), cleared using BABB (benzyl alcohol (#1.09626.1000, Supelco, USA): benzyl benzoate (#10654752, Thermo Fisher Scientific, USA (1:2)) and imaged as follows: Zoom factor 1.6, Ex: 665/45 nm, Em: 725/50 nm (exposure time: 3000 ms (TBEV) and 5000 ms (LGTV and LGTV^T:prME^)) and Ex: 425/60 nm, Em: 480 nm (exposure time: 500 ms), respectively. Pixel intensity range of all images were adjusted and a contrast-limited adaptive histogram equalization (CLAHE) [46] algorithm with a tile size of 8x8 (TBEV and LGTV) or 16x16 (LGTV^T:prME^) was applied. Reconstruction was performed using NRecon software (v.1.7.0.4, Skyscan microCT, Bruker, Belgium) with misalignment compensation and ring artifact reduction. Images were created using Imaris software (v10.1, Bitplane, UK) (viral signal was adjusted to display at min = 7000, max = 30000, and gamma = 1).

### Creation of OPT-MRI fusion images

OPT images with viral signal and autofluorescence signal were reconstructed in DICOM format using NRecon software (v.1.7.0.4, Bruker) followed by their conversion into NifTi using PMOD VIEW tool (v.4.2, PMOD Technologies Inc., Switzerland). OPT-template fusion images were created using the previously described workflow [26]. Briefly, coregistration and normalization of OPT images to the OCUM MRI template was performed using the toolbox SPMmouse in SPM8. All transformation matrices were calculated using individual tissue autofluorescence from OPT images and applied to the corresponding OPT images of the viral staining. Fusion images of viral OPT signal were created for each individual brain using the OCUM template in the PMOD VIEW tool.

### Atlas-based OPT quantification

To assess the differences in viral brain distribution between LGTV, LGTV^T:prME,^ TBEV in more detail and, to provide a metric for viral infection in distinct brain regions, a Volume of Interest (VOI)-based analysis was performed using VOI analysis in the PMOD VIEW tool. Using the OCUM atlas [26], 336 VOIs were automatically delineated, and individual VOI metrics were calculated. Individual thresholded VOI highest pixel intensities were background corrected and all individual VOI metrics of all individual images (LGTV, LGTV^T:prME^ and TBEV) were normalized to their respective median mock intensity in that VOI to obtain an arbitrary unit relative to mock. Thereafter, larger bilateral composite regions of interest (ROIs) were created and the mean value of the arbitrary units of all individual VOIs in a composite ROI were calculated to provide the arbitrary unit in the composite ROI. Finally, ordinary one-way ANOVA with Tukey’s multiple comparison test was performed in GraphPad Prism (v9.3.1 GraphPad Software Inc., San Diego, CA, USA).

### Immunohistochemistry (IHC)

Immunohistochemistry analysis was performed on transverse brain sections from infected or mock-treated mice. Dissection and fixation were carried out as for OPT analysis. Formaldehyde fixed brains were washed 3 times with PBS and dehydrated in a 30% sucrose solution until saturation. Whole brains were snap frozen on dry ice in Optimal Cutting Temperature medium (VWR) and sectioned at 10 μm thick sections on a rotatory microtome cryostat (Microm Microtome Cryostat HM 500M). For staining, sections were permeabilized and blocked (PBS with 10% goat serum, 0.2% TritonX-100, and 1% BSA) for 1 h at room temperature, immunolabelled with primary antibodies against NS5 (chicken polyclonal IgY, dilution 1:1000) and Iba-1 (rabbit polyclonal, Histolab/Biocare CP290, dilution 1:500) or Calbindin C (rabbit monoclonal IgG, Abcam ab108404, 1:500) overnight at 4°C, and then labeled with fluorescent secondary antibody (goat anti-chicken IgY Alexa Fluor™ 555 1:1000 and donkey anti-rabbit IgG Alexa Fluor™ 488 1:1000 Invitrogen A-21206) for 1 h at room temperature. Antibodies were diluted in PBS with 2% goat serum and 0.5% TritonX-100. Confocal laser scanning microscopy was performed using a Leica SP8 confocal microscope and image processing was performed using Imaris Software (v10.1 Bitplane, UK) and ImageJ (NIH).

### Ethics statement

Animal experiments with LGTV and LGTV^T:prME^ were conducted at the Umeå Centre for Comparative Biology (UCCB) and approved by the regional Animal Research Ethics Committee of Northern Norrland and by the Swedish Board of Agriculture under ethical permit A41-2019 and A8-2023. Experiments with TBEV were conducted at an accredited Animal BSL-3 facility at the Veterinary Research Institute, Brno, Czech Republic. The experiments were performed in accordance with Czech laws and guidelines for the use of laboratory animals. The protocol was approved by the Departmental Expert Committee for the Approval of Projects of Experiments on Animals of the Ministry of Agriculture of the Czech Republic and the Committee on the Ethics of Animal Experimentation at the Veterinary Research Institute, Brno, Czech Republic (Approval No. 26674/2020-MZE-18134). For the serological samples, research was carried out in line with “The Code of Ethics of the World Medical Association (Declaration of Helsinki) and according to good clinical practice guidelines. In accordance with local legislation, no formal approval by a research ethics committee was needed, because either anonymous samples or sera for research purposes were used.

### Data availability

The code and scripts used for sequence analysis can be found at https://github.com/Clinical-Genomics-Umea/TBE_analysis. CryoEM density maps have been deposited in the Electron Microscopy Data Bank under EMD accession codes EMD-50624 (Langat), EMD-50518 (LGTV^T:prME^). Atomic coordinates of models have been deposited to the Protein Data Bank (PDB) under accession codes 9FOJ (Langat), 9FK0 and 9H28 (LGTV^T:prME^).

## Additional information

### Author contributions

AKÖ and ER conceptualized the idea and designed the experiments. ER, ML and AM designed, performed molecular cloning and rescued recombinant viruses. ER performed *in vitro* infection experiments. ER, EN, SMAW, AL, JH, PS and DR planned and performed animal experiments. EN, SMAW and ER performed OPT imaging and data analysis. ER, AL and EN performed confocal microscopy. KB, MA and SJB planned the cryo-EM and image reconstruction, purified the virus, acquired data, analyzed the data, prepared figures and deposited the datasets. ES, EM and RN planned the mass spectrometry experiment and analyzed the data. WR performed samples preparation, sequencing and analysis. RN, DR, UA, MA, SJB, EN and AKÖ supervised the work. ER, KB, SMAW and AKÖ wrote the manuscript, and all authors revised the final manuscript.

## Acknowledgments

We would like to thank Gerhard Dobler for kindly providing serum samples from individuals vaccinated against TBEV. We would like to acknowledge The Swedish National Infrastructure (BioMS) and SciLifeLab and their support to the Proteomics Core Facility at Sahlgrenska Academy, University of Gothenburg, for their help with mass spectrometry analysis. Behnam Lak, Pasi Laurinmäki and Ausra Domanska are thanked for their assistance. The facilities and expertise of the HiLIFE CryoEM unit at the University of Helsinki, a member of Instruct-ERIC Centre Finland, FIN-Struct, and Biocenter Finland are gratefully acknowledged. We also acknowledge the Biochemical Imaging Center at Umeå University (BICU), and the National Microscopy Infrastructure for microscopy support (NMI; VR-RFI 2019-00217). The illustration in Figure 1A was prepared using BioRender.com. Some of this research was conducted while SJB visited the Okinawa Institute of Science and Technology through the Theoretical Sciences Visiting Program.

