## Supplementary Material for "The influence of the pre-membrane and envelope proteins on structure, pathogenicity and tropism of tick-borne encephalitis virus"

^1^ Department of Clinical Microbiology. Section of Virology, Umeå University, Umeå, Sweden.

^2^ The Laboratory for Molecular Infection Medicine Sweden (MIMS), Umeå, Sweden.

^3^ Umeå Center for Microbial research. Umeå, Sweden

^4^ Faculty of Biological and Environmental Sciences, Molecular and Integrative Bioscience Research Programme, Helsinki, Finland

^5^ Helsinki Institute of Life Sciences-Institute of Biotechnology, University of Helsinki, Helsinki, Finland.

^6^ Laboratory of Emerging Viral Infections, Veterinary Research Institute, Brno, Czech Republic.

^7^ Faculty of Science, Masaryk University, Brno, Czech Republic.

^8^ Laboratory of Arbovirology, Institute of Parasitology, Biology Centre of the Czech Academy of Sciences, Ceske Budejovice, Czech Republic.

^9^ Faculty of Veterinary Medicine, University of Veterinary Sciences Brno, Brno, Czech Republic.

^10^ Department of Infectious Diseases, Institute of Biomedicine at the Sahlgrenska Academy, University of Gothenburg, Gothenburg, Sweden.

^11^ Department of Clinical Microbiology, Sahlgrenska University Hospital, Västra götalandsregionen, Gothenburg, Sweden.

^12^ Proteomics Core Facility, Sahlgrenska Academy, University of Gothenburg, Gothenburg, Sweden

^13^ Department of Medical and Translational Biology; Umeå University, Umeå, Sweden

^14^ Department of Medical Biosciences, Umeå University, Umeå, Sweden

^15^ Okinawa Institute of Science and Technology, Okinawa, Japan

^16^ Institute of Bioengineering, University of Tartu, Tartu, Estonia.

^$^ Present Address: Institute of Organic Chemistry and Biochemistry, Czech Academy of Sciences, Prague, Czech Republic

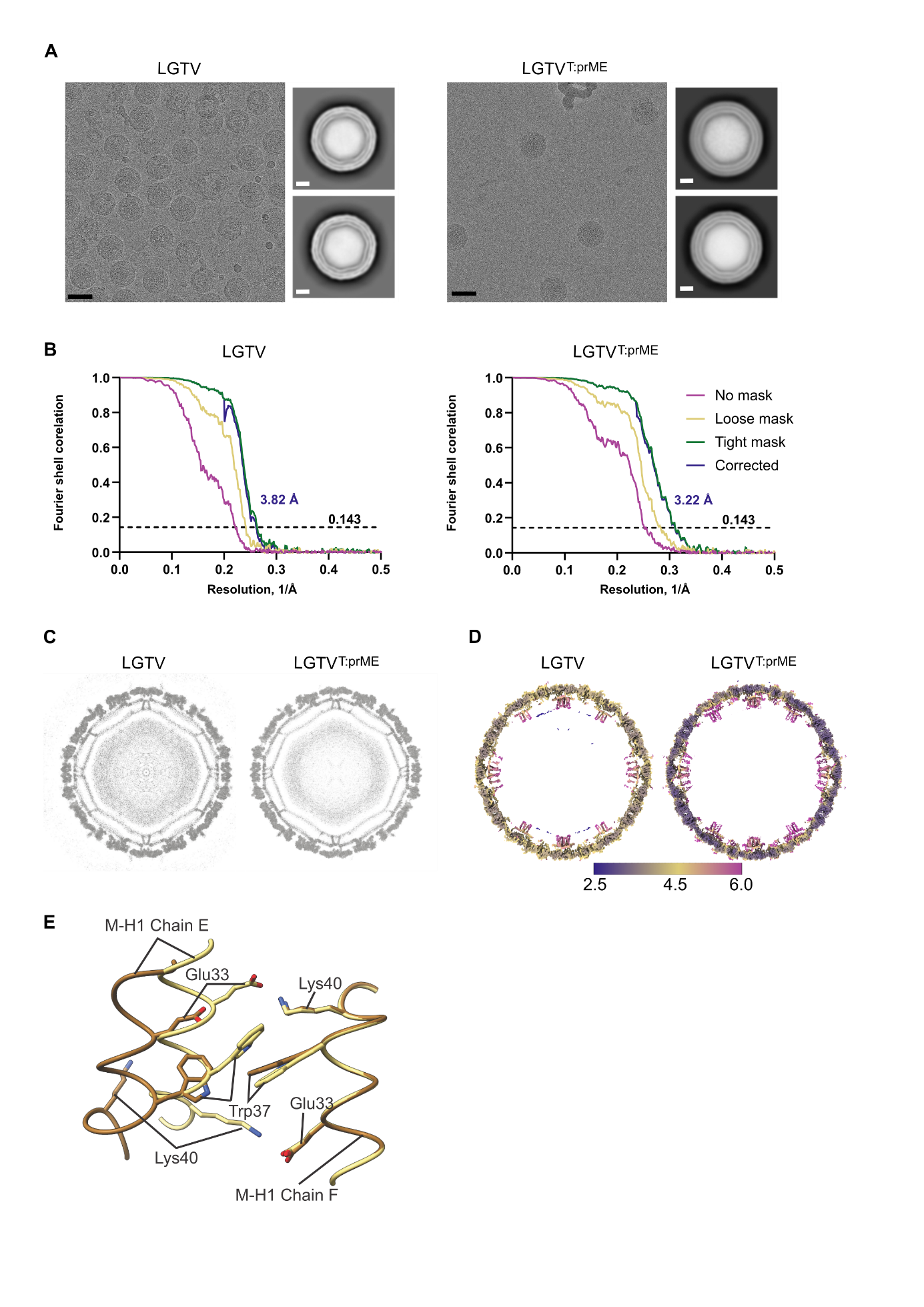

**Supplementary Figure 1. Characterization of LGTV and LGTV^T:prME^ by cryo EM and image reconstruction.** **A)** Representative micrographs of LGTV and chimeric LGTV^T:prME^*,* with the two-dimensional classes picked for reconstruction shown to the right. Scale bar 50 nm (black) or 10 nm (white). **B)** Gold standard Fourier shell correlations LGTV and LGTV^T:prME^ and respectively*.* **C)** Grey scale tilted slab representation through the center of the LGTV reconstruction, 9.69 Å thick, or through LGTV^T:prME^ reconstruction 9.44Å thick. **D)** 30 Å central slices of the LGTV and LGTV^T:prME^ reconstructions, respectively. Viewed down a two-fold axis of symmetry. Colored according to the local resolution, color key in Å.  **E)**  Close-up view of the interaction surface between Chain E and Chain F of the M protein, located between the rafts in LGTV^T:prME^. Conformation consistent with LGTV TP21 (9FOJ) and TBEV Kuutsalo-14 (7Z51) [24] is shown in yellow (deposited as 9FK0). Alternative conformation of Chain E is shown in brown (deposited as 9H28).

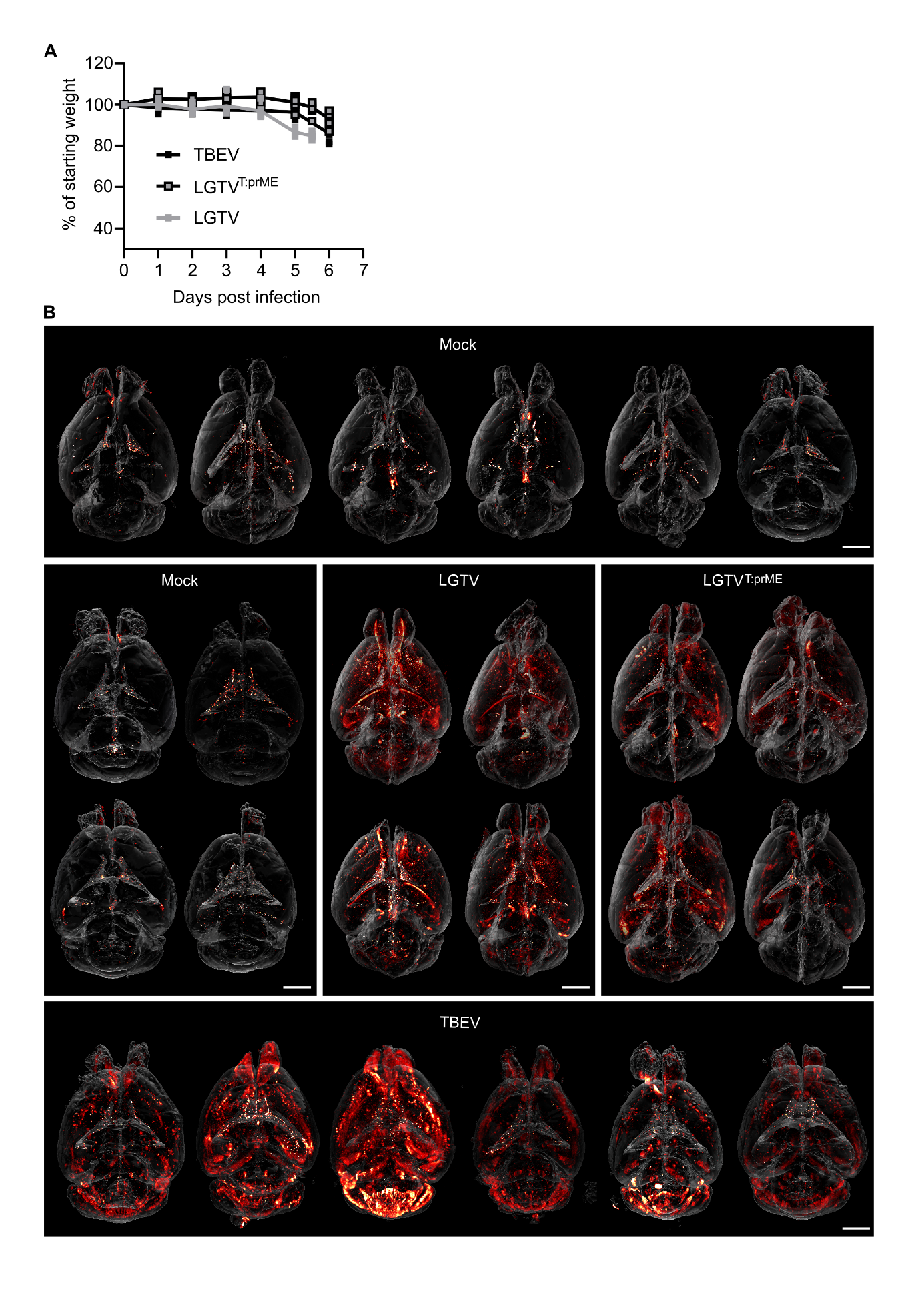

**Supplementary Figure 2. Disease progression and virus infection in whole mouse brain. A)** Weight curves of C57Bl/6 WT mice infected i.c. with LGTV (10^4^ FFU), LGTV^T:prME^ (10^4^ FFU) or TBEV (10^2^ FFU). **B)** Maximum intensity projections (MIP) of virus infection in whole mouse brain visualized by OPT. Brains stained against viral NS5 protein (red). Scale bar 2 mm.

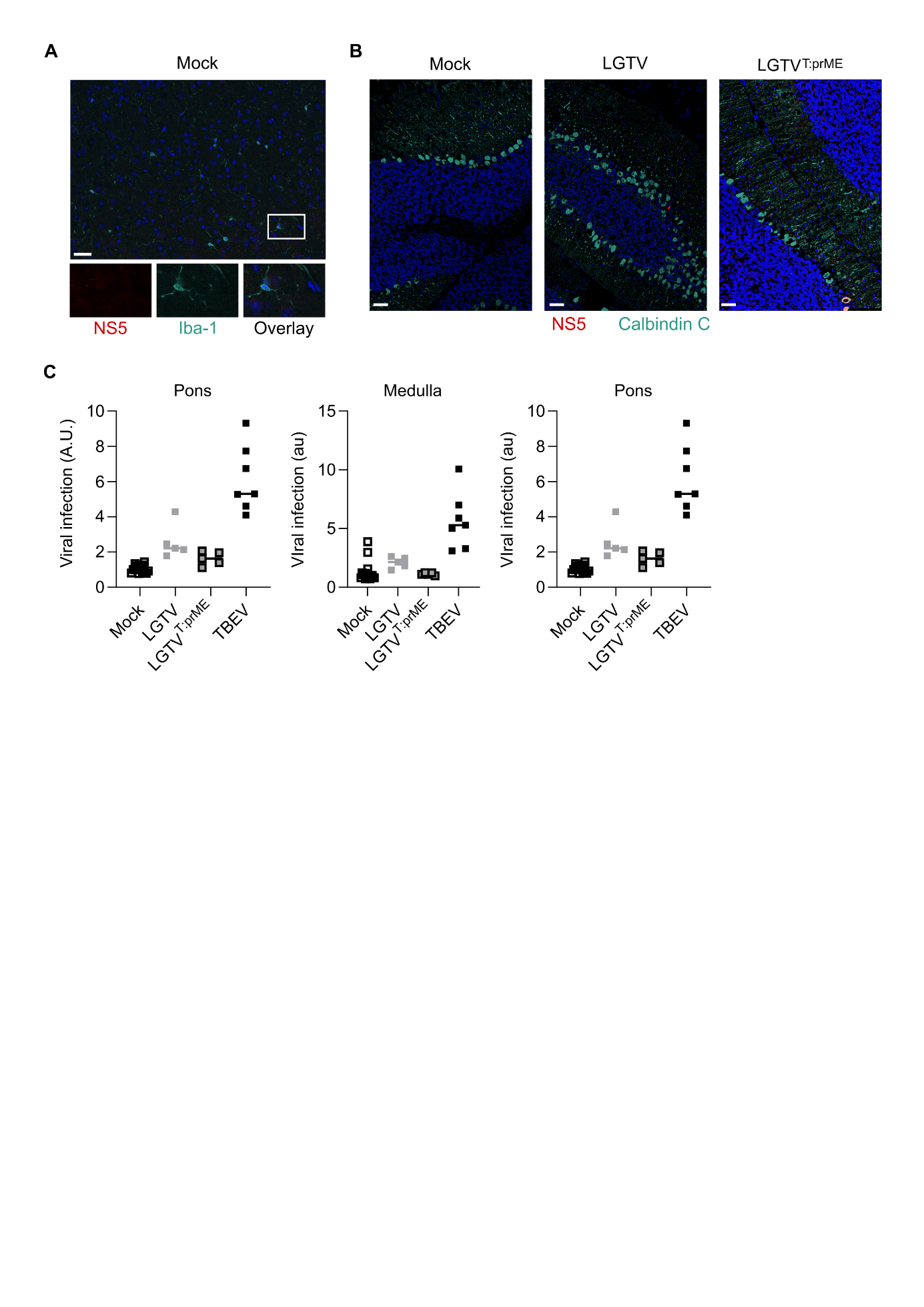

**Supplementary Figure 3. Virus infection in brain by immunohistochemistry and whole brain imaging. A)** Axial brain sections of a mock treated mouse, at the rhinal cortex. Stained against viral NS5 (red) and Iba-1 (turquoise), nuclei are stained with DAPI (blue). **B)** Axial brain sections of cerebellum from a mock treated or infected mice (10^4^ FFU). Stained against viral NS5 (red) and Calbindin C (turquoise), nuclei are stained with DAPI (blue). Images taken by confocal microscopy, magnification 40x. Scale bar 50 μm. **C)** Quantification of viral OPT signal in selected regions of interests (VOIs).

**Supplementary Table 1.** **Mutations in serial passaged LGTV and LGTV^T:prME^ chimeric virus.** Number of mutations present in more than 10% or 50% of all sequencing reads. Result for passage 1 - original stock, and three separate replicates of passage 10. Number of missense mutations, i.e. mutations resulting in an amino acid change, given in parenthesis after total number of mutations.

|  | **Number of mutations**  Total (Missense) | |
| --- | --- | --- |
|  | >10% of all reads | >50% of all reads |
| LGTV passage 1 | 28 (14) | 0 |
| LGTV passage 10.1 | 33 (20) | 3 (3) |
| LGTV passage 10.2 | 32 (18) | 2 (2) |
| LGTV passage 10.3 | 33 (18) | 0 |
| LGTV^T:prME^ passage 1 | 14 (9) | 0 |
| LGTV^T:prME^ passage 10.1 | 15 (10) | 1 (1) |
| LGTV^T:prME^ passage 10.2 | 15 (10) | 2 (2) |
| LGTV^T:prME^ passage 10.3 | 20 (13) | 2 (2) |

**Supplementary Table 2.** **Missense mutations in serial passaged LGTV and chimeric LGTV^T:prME^.** List of all mutations present in majority (>50% of all reads) in at least one replica of the passage 10, together with the resulting amino acid change. Mutations and position given relative to the reference sequence for LGTV TP21 (NCBI reference sequence: NC_003690.1). For each mutation, the percentage of reads with this mutation in passage 1 (original stock) and all three replicates of passage 10 (p10.1 to p10.3) is included.

| **Virus** | **Position (nt)** | **Mutation** | **Affected protein and amino acid substitution** | **Virus passage** | **% of reads**  **with mutation** |
| --- | --- | --- | --- | --- | --- |
| LGTV | 1433 | G->A | E-protein, Glu_153_ ->Lys | p1  **p10.1**  **p10.2**  p10.3 | 2%  **69%**  **85%**  32% |
| LGTV | 9596 | G->A | NS5, Glu_643_->Lys | p1  **p10.1**  **p10.2**  p10.3 | 2%  **56%**  **66%**  28% |
| LGTV | 9734 | G->A | NS5, Asp_691_->Asn | p1  **p10.1**  p10.2  p10.3 | 1%  **57%**  1%  1% |
| LGTV^T:prME^ | 1661 | G->A | E-protein, Gly_231_->Arg | p1  p10.1  **p10.2**  p10.3 | 1%  0%  **65%**  17% |
| LGTV^T:prME^ | 1667 | C->A | E-protein, Gln_233_->Lys | p1  **p10.1**  p10.2  **p10.3** | 1%  **64%**  0%  **61%** |
| LGTV^T:prME^ | 7751 | G->A | NS5, Ala_30_->Thr | p1  p10.1  **p10.2**  p10.3 | 1%  4%  **82%**  18% |
| LGTV^T:prME^ | 9559 | G->T | NS5, Glu_632_->Asp | p1  p10.1  p10.2  **p10.3** | 1%  1%  1%  **71%** |

**Supplementary Table 3.** **Cryo-EM data collection, refinement and validation statistics.**

|  | **LGTV** | Classic conformation  **LGTV^T:prME^** | Alternative conformation  **LGTV^T:prME^** |
| --- | --- | --- | --- |
| Data collection and processing |  |  |  |
| Grid type (carbon) | Quantifoil R2/1.2  2nm continuous carbon | Quantifoil R2/1.2  2nm continuous carbon | Quantifoil R2/1.2  2nm continuous carbon |
| Magnification | 150000 | 150000 | 150000 |
| Voltage (kV) | 200 | 300 | 300 |
| Electron exposure (e–/Å^2^) | 40 | 40 | 40 |
| Defocus range settings (μm) | -0.6 to -1.6 | -0.5 to -1.5 | -0.5 to -1.5 |
| Pixel size (Å) | 0.97 | 0.917 | 0.917 |
| Symmetry imposed | I2 | I2 | I2 |
| Micrographs (no.) | 1430 | 32122 | 32122 |
| Initial particle images (no.) | 15306 | 97855 | 97855 |
| Final particle images (no.) | 12857 | 18724 | 18724 |
| Map resolution (Å)      FSC threshold | 3.82  0.143 | 3.22  0.143 | 3.22  0.143 |
| Map resolution range (Å) | 999 – 1.94 | 999 – 1.83 | 999 – 1.83 |
| Refinement |  |  |  |
| Model composition      Non-hydrogen atoms      Protein residues | 13269  1722 | 13287  1725 | 13287  1725 |
| E | 1-496 | 1-496 | 1-496 |
| M | 1-74 | 1-75 | 1-75 |
| Ligands | NAG, CPL | NAG, CPL | NAG, CPL |
| R.m.s. deviations      Bond lengths (Å)      Bond angles (°) | 0.013  1.848 | 0.012  1.920 | 0.012  1.936 |
| Validation |  |  |  |
| MolProbity score | 0.81 | 0.87 | 0.82 |
| Clashscore | 0.11 | 0.22 | 0.15 |
| Rotamer outliers (%) | 0.29 | 0.57 | 0.86 |
| Ramachandran plot |  |  |  |
| Favoured (%) | 96.17 | 95.94 | 96.12 |
| Allowed (%) | 3.83 | 4.06 | 3.88 |
| Disallowed (%) | 0.00 | 0.00 | 0.00 |

**Supplementary Table 4. Glycosylation of E protein of LGTV^TprME^.** OM = oligomannose.

| **Peptide sequence** | **Glycan Composition** | **# PSMs** | **Theo. MH+ [Da]** | **Precurson ion Abundance (AU)** | **Abundance (% of total observed)** | **Assigned glycan type** |
| --- | --- | --- | --- | --- | --- | --- |
| [Y].TVKVEPHTGDYVAANETHSGRKTASF.[T] | HexNAc(2)Hex(5) | 32 | 4018,80341 | 8,18E+08 | 3,39 | OM |
| [Y].TVKVEPHTGDYVAANETHSGRKTASF.[T] | HexNAc(2)Hex(6) | 62 | 4180,85623 | 4,45E+09 | 18,43 | OM |
| [Y].TVKVEPHTGDYVAANETHSGRKTASF.[T] | HexNAc(2)Hex(7) | 49 | 4342,90905 | 4,23E+09 | 17,53 | OM |
| [Y].TVKVEPHTGDYVAANETHSGRKTASF.[T] | HexNAc(2)Hex(8) | 41 | 4504,96188 | 3,46E+09 | 14,34 | OM |
| [Y].TVKVEPHTGDYVAANETHSGRKTASF.[T] | HexNAc(2)Hex(9) | 17 | 4667,0147 | 5,26E+08 | 2,18 | OM |
| [Y].TVKVEPHTGDYVAANETHSGRKTASF.[T] | HexNAc(3)Hex(4) | 18 | 4059,82996 | 3,06E+08 | 1,27 | hybrid |
| [Y].TVKVEPHTGDYVAANETHSGRKTASF.[T] | HexNAc(3)Hex(4)Fuc(1) | 23 | 4205,88787 | 1,11E+09 | 4,60 | hybrid |
| [Y].TVKVEPHTGDYVAANETHSGRKTASF.[T] | HexNAc(3)Hex(4)Fuc(1)NeuAc(1) | 15 | 4496,98328 | 4,90E+08 | 2,03 | hybrid |
| [Y].TVKVEPHTGDYVAANETHSGRKTASF.[T] | HexNAc(3)Hex(5) | 22 | 4221,88278 | 6,40E+08 | 2,65 | hybrid |
| [Y].TVKVEPHTGDYVAANETHSGRKTASF.[T] | HexNAc(3)Hex(5)Fuc(1) | 28 | 4367,94069 | 9,98E+08 | 4,13 | hybrid |
| [Y].TVKVEPHTGDYVAANETHSGRKTASF.[T] | HexNAc(3)Hex(5)Fuc(1)NeuAc(1) | 12 | 4659,03611 | 4,15E+08 | 1,72 | hybrid |
| [Y].TVKVEPHTGDYVAANETHSGRKTASF.[T] | HexNAc(3)Hex(5)NeuAc(1) | 22 | 4512,9782 | 2,63E+08 | 1,09 | hybrid |
| [Y].TVKVEPHTGDYVAANETHSGRKTASF.[T] | HexNAc(3)Hex(6) | 39 | 4383,9356 | 1,35E+09 | 5,60 | hybrid |
| [Y].TVKVEPHTGDYVAANETHSGRKTASF.[T] | HexNAc(3)Hex(6)Fuc(1) | 28 | 4529,99351 | 3,01E+08 | 1,25 | hybrid |
| [Y].TVKVEPHTGDYVAANETHSGRKTASF.[T] | HexNAc(3)Hex(6)NeuAc(1) | 19 | 4675,03102 | 6,02E+08 | 2,49 | hybrid |
| [Y].TVKVEPHTGDYVAANETHSGRKTASF.[T] | HexNAc(4)Hex(5) | 21 | 4424,96215 | 1,64E+08 | 0,68 | complex |
| [Y].TVKVEPHTGDYVAANETHSGRKTASF.[T] | HexNAc(4)Hex(5)Fuc(1) | 40 | 4571,02006 | 2,54E+09 | 10,51 | complex |
| [Y].TVKVEPHTGDYVAANETHSGRKTASF.[T] | HexNAc(4)Hex(5)Fuc(1)NeuAc(1) | 31 | 4862,11548 | 1,25E+09 | 5,19 | complex |
| [Y].TVKVEPHTGDYVAANETHSGRKTASF.[T] | HexNAc(4)Hex(5)Fuc(1)NeuAc(2) | 10 | 5153,21089 | 2,23E+08 | 0,92 | complex |

**Supplementary Table 5. Viral infection metrics of the 20 highest infected brain regions from atlas-based quantification.** For TBEV, all cerebellar regions were taken out of this analysis to compare the highest infected regions in cerebrum and OB.

| **LGTV** |  |  |  |  |  |  |  |  |
| --- | --- | --- | --- | --- | --- | --- | --- | --- |
|  | **A** | **B** | **C** | **D** | **E** |  |  | **Mean** |
| Frontal association cortex | 3.9 | 3.4 | 4.5 | 2.2 | 3.0 |  |  | 3.4 |
| Caudomedial entorhinal cortex | 4.0 | 4.4 | 2.3 | 3.4 | 2.1 |  |  | 3.2 |
| Anterior olfactory  Nucleus | 4.3 | 1.6 | 4.0 | 2.6 | 3.3 |  |  | 3.2 |
| Piriform cortex | 2.9 | 2.3 | 2.9 | 2.3 | 4.8 |  |  | 3.1 |
| Olfactory bulb:  Granule cell layer | 3.2 | 2.6 | 3.5 | 1.9 | 4.0 |  |  | 3.0 |
| Perirhinal cortex | 4.8 | 3.0 | 1.5 | 2.7 | 2.7 |  |  | 2.9 |
| Cingulate cortex, area 32 | 3.9 | 2.6 | 2.4 | 2.6 | 3.1 |  |  | 2.9 |
| Primary somatosensory cortex | 4.4 | 2.2 | 1.8 | 2.8 | 3.3 |  |  | 2.9 |
| Medial orbital cortex | 4.0 | 2.5 | 2.4 | 2.3 | 2.9 |  |  | 2.8 |
| Primary somatosensory cortex: hindlimd | 3.6 | 2.2 | 1.5 | 2.5 | 4.3 |  |  | 2.8 |
| Anterior commissure:  Olfactory limb | 3.3 | 2.2 | 3.1 | 1.4 | 4.1 |  |  | 2.8 |
| Hippocampal region  MoDG | 3.3 | 2.3 | 2.8 | 2.6 | 3.0 |  |  | 2.8 |
| Insular region | 4.5 | 2.1 | 2.2 | 2.3 | 2.8 |  |  | 2.8 |
| Olfactory Tubercle | 2.9 | 2.9 | 2.1 | 2.3 | 3.2 |  |  | 2.7 |
| Frontal cortex: area 3 | 2.8 | 1.6 | 2.4 | 2.5 | 3.9 |  |  | 2.7 |
| Secondary motor cortex | 3.5 | 2.7 | 2.3 | 2.1 | 2.6 |  |  | 2.7 |
| Olfactory peduncle | 3.3 | 2.1 | 2.3 | 2.0 | 3.5 |  |  | 2.6 |
| Lateral Olfactory tract | 2.8 | 2.0 | 3.7 | 1.4 | 3.3 |  |  | 2.6 |
| Cerebellar vermis lob1-2 | 4.0 | 1.7 | 2.6 | 2.8 | 1.9 |  |  | 2.6 |
| Hippocampal region: GrDG | 2.4 | 1.7 | 2.6 | 3.4 | 2.8 |  |  | 2.6 |
| **LGTV^T:prME^** |  |  |  |  |  |  |  |  |
|  | **F** | **G** | **H** | **I** | **J** |  |  | **Mean** |
| Cingulate cortex, area 32 | 2.6 | 2.4 | 4.3 | 2.8 | 1.9 |  |  | 2.8 |
| Dorsolateral entorhinal  cortex | 3.3 | 2.3 | 4.4 | 1.8 | 2.0 |  |  | 2.8 |
| Medial entorhinal cortex | 3.2 | 2.5 | 3.6 | 1.7 | 1.8 |  |  | 2.5 |
| Caudomedial entorhinal cortex | 3.1 | 2.5 | 3.6 | 1.7 | 1.8 |  |  | 2.6 |
| Perirhinal cortex | 2.6 | 2.1 | 4.0 | 1.9 | 1.7 |  |  | 2.4 |
| Cerebral peduncle | 1.7 | 1.5 | 4.2 | 2.8 | 1.7 |  |  | 2.4 |
| Amygdala | 2.6 | 1.7 | 3.6 | 1.8 | 1.9 |  |  | 2.3 |
| Ectorhinal cortex | 2.2 | 2.4 | 3.4 | 2.1 | 1.5 |  |  | 2.3 |
| Temporal Association Area | 2.0 | 1.9 | 2.9 | 3.1 | 1.6 |  |  | 2.3 |
| Cerebellar vermis lob1-2 | 2.6 | 2.9 | 2.3 | 1.2 | 2.1 |  |  | 2.2 |
| Primary Visual cortex: monocular area | 1.7 | 1.7 | 2.0 | 4.3 | 1.4 |  |  | 2.2 |
| Hippocampal region PoDG | 1.6 | 1.4 | 5.2 | 1.9 | 1.2 |  |  | 2.2 |
| Ventral claustrum | 1.3 | 2.1 | 4.6 | 1.8 | 1.2 |  |  | 2.2 |
| Primary somatosensory cortex: hindlimd | 2.0 | 1.7 | 2.4 | 3.1 | 1.4 |  |  | 2.1 |
| Secondary visual cortex: lateral area | 2.0 | 2.2 | 2.8 | 1.8 | 1.6 |  |  | 2.1 |
| Insular region | 1.9 | 2.7 | 3.1 | 1.5 | 1.2 |  |  | 2.1 |
| Frontal association cortex | 1.6 | 3.0 | 2.9 | 1.5 | 1.4 |  |  | 2.1 |
| Medial orbital cortex | 2.1 | 2.0 | 2.4 | 2.4 | 1.4 |  |  | 2.1 |
| Primary motor cortex | 1.9 | 1.8 | 3.9 | 1.4 | 1.2 |  |  | 2.0 |
| Anterior commissure:  Olfactory limb | 3.0 | 1.4 | 2.8 | 1.3 | 1.3 |  |  | 2.0 |
| **TBEV** |  |  |  |  |  |  |  |  |
|  | **K** | **L** | **M** | **N** | **O** | **P** | **Q** | **Mean** |
| Piriform cortex | 11.7 | 6.1 | 6.7 | 6.2 | 8.4 | 5.3 | 4.6 | 7.0 |
| Pons | 7.7 | 5.3 | 6.7 | 4.1 | 9.3 | 4.6 | 5.3 | 6.1 |
| Medulla | 7.0 | 5.9 | 5.3 | 5.0 | 10.1 | 3.3 | 3.1 | 5.7 |
| Ventral Tenia Tecta | 10.4 | 6.0 | 4.8 | 6.3 | 6.3 | 1.8 | 3.5 | 5.6 |
| Olfactory peduncle | 10.0 | 4.1 | 5.3 | 5.2 | 6.6 | 2.7 | 4.1 | 5.4 |
| Insular region | 7.9 | 3.9 | 6.7 | 4.5 | 6.7 | 4.1 | 3.8 | 5.4 |
| Primary somatosensory cortex: Barrel field | 7.0 | 2.8 | 6.6 | 5.0 | 9.2 | 3.3 | 2.9 | 5.3 |
| Caudomedial entorhinal cortex | 6.6 | 4.7 | 4.9 | 5.3 | 8.0 | 3.8 | 3.5 | 5.3 |
| Ventral Intermediate  Entorhinal cortex | 6.8 | 4.2 | 4.1 | 4.3 | 10.1 | 3.8 | 4.0 | 5.3 |
| Secondary motor cortex | 9.4 | 4.7 | 5.1 | 4.1 | 5.9 | 4.0 | 3.5 | 5.2 |
| Medial entorhinal cortex | 7.2 | 4.7 | 5.1 | 4.5 | 7.3 | 3.9 | 3.9 | 5.2 |
| Hippocampal region  CA20r | 9.9 | 3.6 | 3.5 | 2.8 | 12.7 | 1.2 | 2.0 | 5.1 |
| Posteromedial cortical  Amygdaloid area | 6.9 | 6.3 | 4.0 | 4.5 | 7.4 | 3.0 | 3.9 | 5.1 |
| Cingulate cortex: area 30 | 7.4 | 3.8 | 4.2 | 4.7 | 8.1 | 3.3 | 3.5 | 5.0 |
| Hippocampal region: Slu | 8.5 | 4.3 | 5.4 | 3.0 | 10.0 | 1.8 | 1.5 | 4.9 |
| Dorsolateral entorhinal  cortex | 7.4 | 5.6 | 4.7 | 4.0 | 7.0 | 3.1 | 2.7 | 4.9 |
| Olfactory tubercle | 7.4 | 4.2 | 5.3 | 4.3 | 6.5 | 2.1 | 4.4 | 4.9 |
| Mamillary bodies | 8.2 | 4.4 | 3.6 | 3.4 | 8.4 | 1.9 | 3.5 | 4.8 |
| Hypothalamus | 8.6 | 5.6 | 2.9 | 5.2 | 5.7 | 2.0 | 3.6 | 4.8 |
| Medial Orbital cortex | 9.4 | 3.5 | 3.8 | 6.0 | 5.0 | 2.7 | 3.1 | 4.8 |
