## Supplementary figures and images for "The influence of the pre-membrane and envelope proteins on structure, pathogenicity and tropism of tick-borne encephalitis virus"

### Supplementary Figure 1

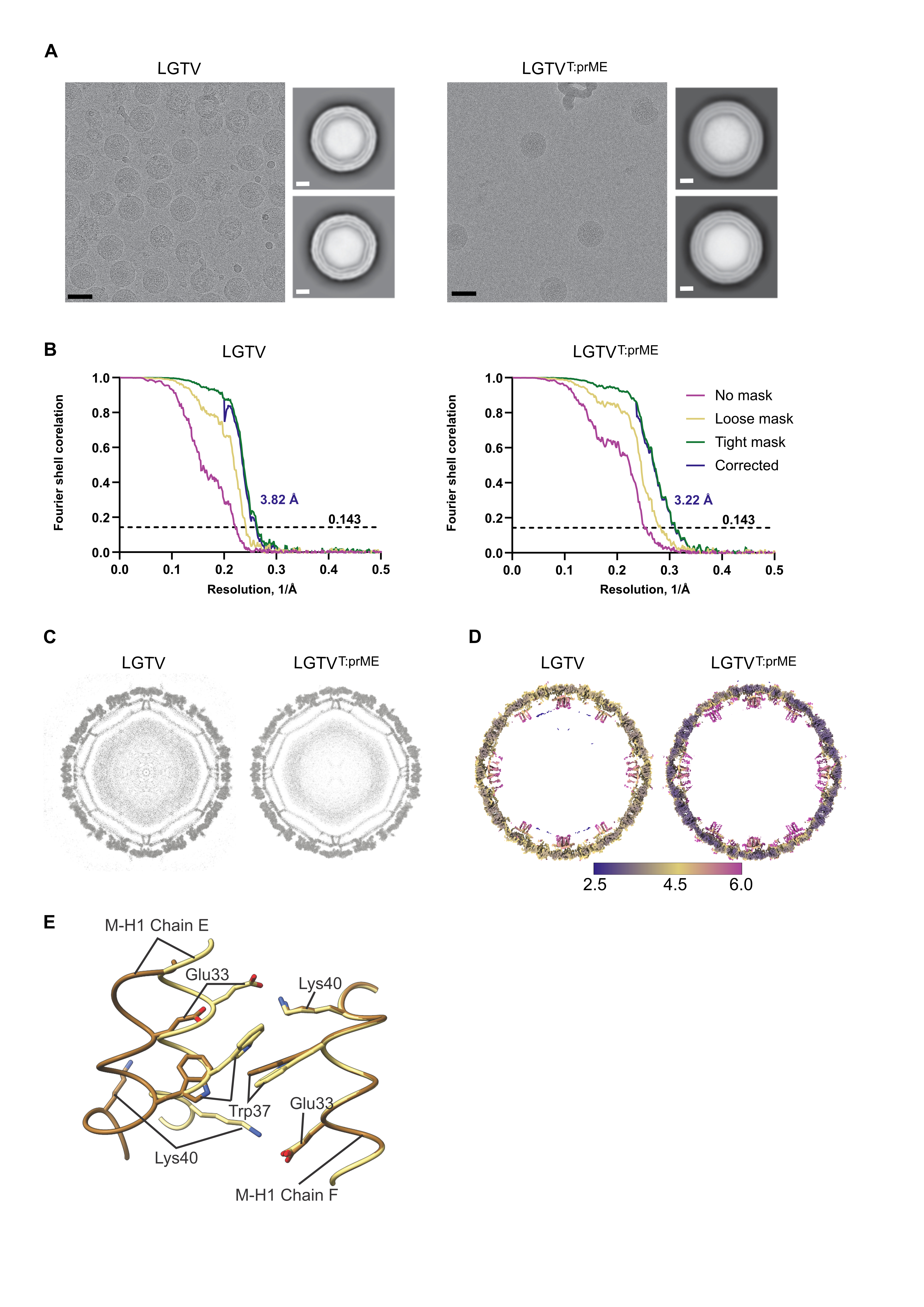

### Supplementary Figure 2

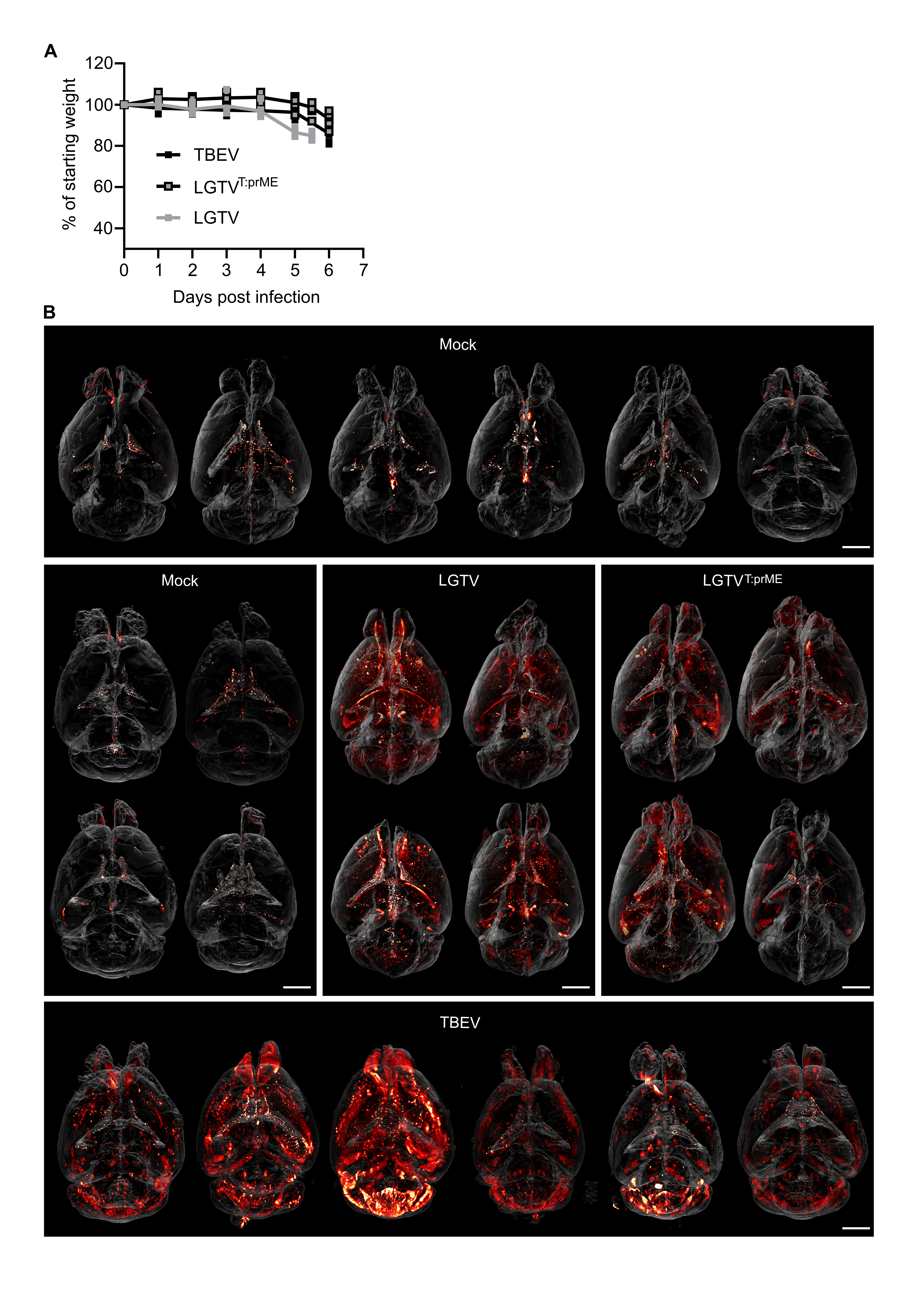

### Supplementary Figure 3

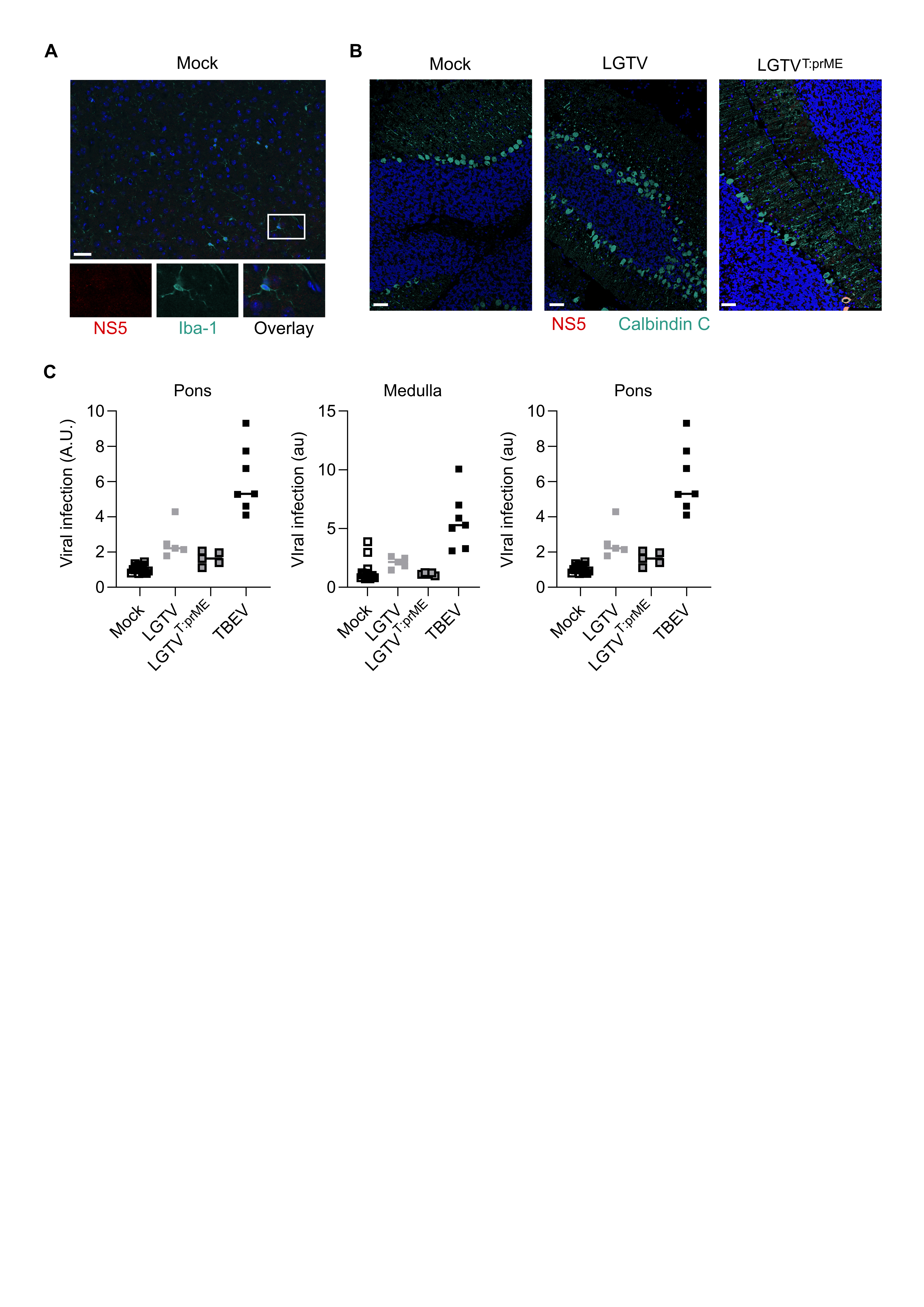
